# Unraveling Tissue-Specific Molecular Signatures and Convergent Pathway Enrichments in Suicidal Behavior

**DOI:** 10.64898/2026.02.27.708508

**Authors:** Aaron K. Jenkins, Meilin Jia-Richards, Madeline R. Scott, Eli Goodfriend, RuoFei Yin, Sarah Riston, Kyle D. Ketchesin, Haeun Moon, Kaitlyn Petersen, Antoine Douaihy, Jill R. Glausier, David A. Brent, David A. Lewis, Anna Marsland, George C. Tseng, Kehui Chen, Marianne L. Seney, Colleen A. McClung, Nadine M. Melhem

## Abstract

Suicide is a leading cause of death worldwide, yet the biological mechanisms underlying suicide remain poorly understood. A clearer understanding at the molecular level is essential for developing objective biomarkers and targeted interventions. In this study, we used transcriptomic profiling to investigate gene expression patterns associated with suicidal thoughts and behaviors across peripheral blood (n=264) and postmortem brain tissue from two prefrontal regions (dorsolateral prefrontal cortex, DLPFC; subgenual anterior cingulate cortex, sgACC) of individuals with and without psychiatric illness (n=249). Peripheral analyses revealed broad transcriptional changes associated with suicidal thoughts and behaviors, marked by dysregulated immune-related and inflammatory processes. Longitudinal modeling further revealed gene co-expression modules that predicted future suicide attempts over a 12-month follow-up, highlighting processes related to apoptosis, mitochondrial function, and immune regulation. By contrast, transcriptomic analyses of postmortem tissue derived from the DLPFC and sgACC revealed largely suppressed neuroimmune activity. Gene co-expression analyses in the brain identified suicide-associated modules enriched for synaptic plasticity, oxidative stress, and neuroimmune function, some of which displayed regional specificity. Cross-tissue comparison showed minimal gene-level overlap between brain and blood, although shared pathway-level themes emerged in immune, sensory, and cellular stress processes. Taken together, these findings suggest that suicide is associated with distinct but functionally convergent transcriptional alterations across brain and blood. By integrating tissue-specific and systems-level molecular signatures, this work provides insight into the biological architecture of suicide and lays the groundwork for developing novel biomarkers and therapeutic targets to improve prevention and treatment outcomes.

## INTRODUCTION

Suicide is a significant public health concern, responsible for nearly 746,000 deaths globally each year^1^. While rates have declined in some regions, suicide continues to rank as the third leading cause of death among individuals aged 15 to 29 worldwide^2^ and the second leading cause among adolescents and young adults in the United States^3^. Suicide is a complex and heterogeneous entity shaped by the confluence of multiple biological, environmental, and psychosocial factors^4,5^. Accurately identifying individuals at risk of suicide and improving prevention efforts remain formidable challenges, largely due to gaps in our understanding of the underlying neurobiological mechanisms.

Evidence from genetic studies supports a heritable component to suicide risk^6^. Genome-wide association studies have begun to identify risk loci associated with suicide,^7^ but these findings offer limited insight into how genetic variation translates into altered molecular function. To bridge this gap, transcriptomic approaches have examined molecular signatures in both postmortem brain tissue and peripheral blood, aiming to capture central neurobiological alterations alongside more accessible biomarkers of suicide risk^8^. By capturing downstream molecular activity, transcriptomic analyses provide a functional readout of gene regulation, offering a window into the biological effects of genetic and environmental risk factors in the brain.

Postmortem transcriptomic studies have been invaluable in identifying biological pathways implicated in suicide^9–12^. However, most prior work has focused on single diagnoses or brain regions, even though suicide transcends diagnostic categories and involves distributed neural circuits related to emotion regulation, impulse control, and cognitive processing^13^. Examining multiple, interconnected cortical regions within transdiagnostic cohorts can provide a more comprehensive view of suicide-related molecular alterations.

To build on insights from postmortem research, longitudinal transcriptomic profiling in living individuals can reveal dynamic molecular changes across the continuum of suicide risk^14,15^. While temporal resolution provides insight into how molecular states evolve within individuals, it does not address how biological signals are distributed across tissues. Integrating cross-tissue approaches therefore adds an orthogonal dimension of insight, enabling comparison between central and peripheral compartments. Such strategies are especially valuable because they challenge the common assumption that molecular alterations observed in peripheral blood necessarily mirror those in the brain. Instead, peripheral and central tissues may exhibit distinct, yet complementary molecular responses to upstream stressors. Examining both compartments is therefore critical for capturing a more comprehensive illustration of the biological processes associated with suicide risk and for informing biologically-grounded models of prevention and intervention^16,17^.

Taken together, these considerations underscore the necessity for integrative approaches. The current study combines longitudinal peripheral transcriptomic data from living individuals with postmortem brain data from suicide decedents and matched comparison groups. By combining these two complementary sources of data, we aim to identify molecular pathways and gene co-expression networks associated with suicidal ideation, suicide attempt, and suicide death. Our postmortem analyses focus on two brain regions critically implicated in the neurobiology of suicide: the dorsolateral prefrontal cortex (DLPFC),^18,19^ which subserves cognitive control, and the subgenual anterior cingulate cortex (sgACC),^20,21^ which mediates emotion regulation. These regions are reciprocally connected, functioning in concert to regulate these processes. This approach leverages a transdiagnostic, multiregional postmortem brain cohort alongside a prospective, clinically characterized peripheral blood cohort. The primary goals of this study are to (1) identify gene expression patterns that predict suicidal behaviors in peripheral blood, (2) characterize transcriptomic signatures of suicide death across the DLPFC and sgACC, and (3) evaluate whether peripheral and central molecular alterations converge or diverge, and how these patterns inform biologically-grounded models of suicide risk.

## MATERIALS AND METHODS

### Subjects

#### Peripheral Cohort

Participants were recruited from the University of Pittsburgh Medical Center (UPMC) Western Psychiatric Hospital and a university-affiliated research registry. Psychiatric patients across the spectrum of psychiatric diagnoses and healthy controls were enrolled. After quality control, gene expression data from 264 participants were included.

Participants were classified as recent suicide attempters (SA, n=69), individuals with suicidal ideation without recent attempt (SI, n=81), psychiatric controls without suicidal thoughts or behaviors (PC, n=73), or healthy controls without psychiatric history (HC, n=41). Unless otherwise specified, SA and SI participants were considered suicidal participants (SP). During longitudinal follow-up, 17 participants made an actual suicide attempt and 36 exhibited suicidal behaviors, including actual, interrupted, or aborted attempts, preparatory behaviors, or emergency referral for suicidal ideation.

#### Postmortem Cohort

Tissue from the DLPFC (BA46) and sgACC (BA25) was collected from subjects with a psychiatric diagnosis (n=166), including major depressive disorder (MDD; n=83), bipolar disorder (BP; n=33), and schizophrenia (SCZ; n=50), and nonpsychiatric comparison subjects (NPC; n=83). Postmortem brain tissue was obtained from the Allegheny County Medical Examiner’s Office with next-of-kin consent and diagnoses were determined via psychological autopsy, medical record review, and DSM-5 consensus^22^. All procedures were approved by the University of Pittsburgh IRB and the Committee for Oversight of Research and Clinical Training Involving Decedents. For our analysis, the subjects with a psychiatric diagnosis were further divided into suicide decedents with psychiatric diagnoses (SP, n=50) and non-suicide psychiatric comparisons (NSP, n=116) (**Table 1**).

**Table 1.** Summary of subject characteristics for the proposed studies. Abbreviations: A, Asian; B, Black; F, female; **M_1_** male; O, other; PMl, postmortem interval; RIN,. RNA integrity number; W, White. Although not included here, we have information about psychotropic medications in each subject.

| Postmortem Brain Subjets |  |  |  |  |
| --- | --- | --- | --- | --- |
|  | SS (n=50) | NSS (n=116) | NPC (n=83) |  |
| Sex, No. | 38 M / 12 F | 77 M / 39 F | 58 M / 25 F |  |
| Race | 43 W / 5 B / 2 A | 95 W / 21 B | 66 W / 17 B |  |
| Age, mean ± SD, y | 41.8 ± 12.5 | 48.8 ± 9.3 | 47.2 ± 12.2 |  |
| PMI, mean ± SD, h | 17.8 ± 6.2 | 17.3 ± 6.2 | 16.9 ± 5.7 |  |
| Brain pH, mean ± SD | 6.7 ± 0.3 | 6.5 ± 0.3 | 6.6 ± 0.3 |  |
| RIN, mean ± SD | 8.2 ± 0.8 | 7.9 ± 0.8 | 8.0 ± 0.8 |  |
| Psychiatric Diagnoses |  |  |  |  |
| MDD | 30 | 53 | 0 |  |
| BP | 9 | 24 | 0 |  |
| SCZ | 11 | 39 | 0 |  |
| Living Peripheral Subjects |  |  |  |  |
|  | SA (n=51) | SI (n=60) | PC (n=42) | HC (n=34) |
| Sex, No. | 24 M / 27 F | 30 M / 29 F / 1 O | 22 M / 20 F | 15 M / 19 F |
| Race | 32W/11B/2A/6O | 48W/4B/5A/3O | 32W/6B/1A/3O | 28W/3B/1A/2O |
| Age, mean ± SD, y | 24 ± 3.7 | 24.7 ± 3.9 | 24.4 ± 3.6 | 24.9 ± 3.6 |
| Psychiatric Diagnoses |  |  |  |  |
| Mood disorder | 44 | 54 | 33 | 0 |
| Anxiety disorder | 17 | 28 | 24 | 0 |
| Psychotic disorder | 2 | 5 | 2 | 0 |
| Trauma disorder | 19 | 14 | 10 | 0 |
| Substance use disorder | 28 | 40 | 17 | 0 |

### Sample Processing, RNA Sequencing, and Gene Expression Preprocessing

#### Peripheral Blood

Whole blood (2.5 mL) was collected at baseline and 3-, 6-, and 12-month follow-ups using PAXgene Blood RNA tubes. RNA was extracted and amplified using the Illumina TotalPrep RNA Amplification Kit (Ambion). RNA concentration and integrity were assessed at extraction, and samples failing quality threshold were excluded.

Gene expression profiling was performed using Affymetrix Clariom S HT microarray platform (Thermo Fisher Scientific). Raw data were processed using the Transcriptome Analysis Console software, where standard quality control metrics (labeling, hybridization, and signal distribution) were evaluated. Gene-level expression data were filtered and normalized for downstream analyses.

#### Postmortem Brain

RNA was extracted from frozen DLPFC and sgACC tissue using TRIzol (Invitrogen) followed by purification with the RNeasy Lipid Tissue Mini Kit (Qiagen) as previously described^23^. RNA quality and quantity were assessed via spectrophotometry (Qubit) and Agilent Bioanalyzer. RNA-seq libraries were prepared using the TruSeq Stranded Total RNA Library Prep Kit (Illumina) with rRNA depletion and sequenced on Illumina NovaSeq 6000 (paired-end, 101 bp; mean depth = 45.7 million reads/sample). Reads were quality checked with FastQC, trimmed using Trimmomatic, and aligned to GRCh38 with HISAT2^24^. Gene-level counts were generated using HTSeq (v0.11.2)^25^.

Gene expression matrices initially contained 60,676 Ensembl genes. Low-expression genes were filtered using edgeR’s *filterByExpr* function, retaining genes with sufficient counts across samples (minimum group size n=33). After filtering across both brain regions, 16,884 genes remained for downstream analyses.

#### Differential Expression

To characterize transcriptomic alterations associated with suicidal thoughts and behaviors in peripheral blood, we evaluated three a priori contrasts: 1) SP vs. HC, 2) SP vs. PC, and 3) SI vs. SA. These contrasts were designed to capture deviations from normative expression, isolate suicide-related differences beyond those related to psychiatric diagnoses and distinguish molecular features of ideation versus attempt.

For postmortem brain analyses, gene expression was examined in the DLPFC and sgACC across three contrasts: 1) SP vs. NPC, 2) SP vs. NSP, and 3) NSP vs. NPC. These comparisons enabled dissociation of suicide-related transcriptional changes from those associated with psychiatric illness alone.

Differential expression (DE) analyses were performed using DESeq2^26^ with group status modeled as the primary factor and relevant covariates included as appropriate. To balance sensitivity and specificity in this exploratory context, genes were considered DE at p<0.05 with an absolute log_2_ fold change > 0.26.

#### Co-expression Network Analyses

Gene co-expression patterns were examined using Weighted Gene Co-expression Network Analysis (WGCNA). Expression matrices were screened using the *goodSamplesGenes function* and soft-thresholding powers were selected to approximate scale-free topology (fit index ≥ 0.9). Adjacency matrices were transformed into topological overlap matrices followed by hierarchical clustering and dynamic tree cutting to identify co-expression modules. Associations between module eigengenes and suicide-related phenotypes were quantified using distance correlation (dCor) with 10,000 permutations, and module robustness was assessed using a frequent-voting stability procedure^27^. Modules were prioritized based on significant associations, permutation support, and stability estimates above 0.20.

In peripheral blood, prospective associations between baseline gene expression and future suicide attempt (SA) or broader suicidal behavior (SB) were evaluated using logistic regression. For longitudinal gene expression, we used Cox proportional hazards models with time-varying covariates. Bonferroni correction was applied to account for testing across the two prospective outcomes.

#### Biological Process Enrichment Analysis

Pathway enrichment analyses were performed separately for blood and brain datasets. In peripheral blood, Gene Set Enrichment Analysis was performed using MSigDB gene sets to identify pathway-level alterations across group comparisons^28,29^. In postmortem brain tissue, DE genes from the DLPFC and sgACC were analyzed using Metascape^30^. Enrichment analyses focused on biological processes and molecular functions, using all expressed genes as the background.

#### Cross-Tissue and Cross-Region Overlap

To assess shared transcriptional patterns across tissues and brain regions, we applied rank-rank hypergeometric overlap (RRHO), a threshold-free method that detects concordance between two ranked gene lists based on effect size and statistical significance^31,32^. RRHO analyses were used to compare transcriptomic profiles between cortical regions (DLPFC and sgACC) and between blood and brain tissue.

To complement this rank-based analyses, we performed set-based overlap analyses to explicitly quantify shared and opposing DE across tissues. DE genes were defined using an exploratory threshold of p<0.05, yielding approximately 12,400 genes available for cross-tissue intersection. Gene sets were stratified by direction of effect and visualized using UpSet plot, enabling direct comparison of same-direction and opposing expression patterns between blood and brain.

## RESULTS

### Immune and Sensory Dysregulation Characterize Suicidal Phenotypes in Blood

To identify peripheral molecular signatures associated with suicidal behavior, we performed DE analyses across three contrasts: (1) suicidal participants (SP) vs. healthy controls (HC), capturing transcriptional differences associated with suicidal ideation or attempt relative to individuals without psychiatric illness; (2) SP vs. psychiatric controls (PC), isolating suicide-associated molecular changes beyond psychiatric diagnoses; and (3) suicide attempt (SA) vs. suicidal ideation (SI), distinguishing individuals who had attempted suicide from those reporting ideation alone. Genes were considered DE at a p-value < 0.05 and an absolute log2 fold change > 0.26, a threshold selected to balance statistical rigor with sensitivity to biologically meaningful transcriptional shifts in peripheral blood.

DE analyses revealed widespread molecular alterations across all comparisons. Substantial differences were observed in SP vs. HC (**Figure 1A**; 1,026 DE genes, 826 upregulated, 200 downregulated), while the largest number of DE genes emerged in the SP vs. PC comparison (**Figure 1B**; 1,061 DE genes, 198 upregulated, 863 downregulated), indicating robust suicide-associated transcriptional differences beyond psychiatric diagnosis. In contrast, SI vs. SA comparison identified fewer DE genes overall (**Figure 1C**; 347 DE genes, 294 upregulated, 53 downregulated). Although fewer genes reached statistical significance in this contrast, distinct clusters of DE transcripts were observed, indicating measurable molecular differences between suicidal ideation and attempt.

**Figure 1.**
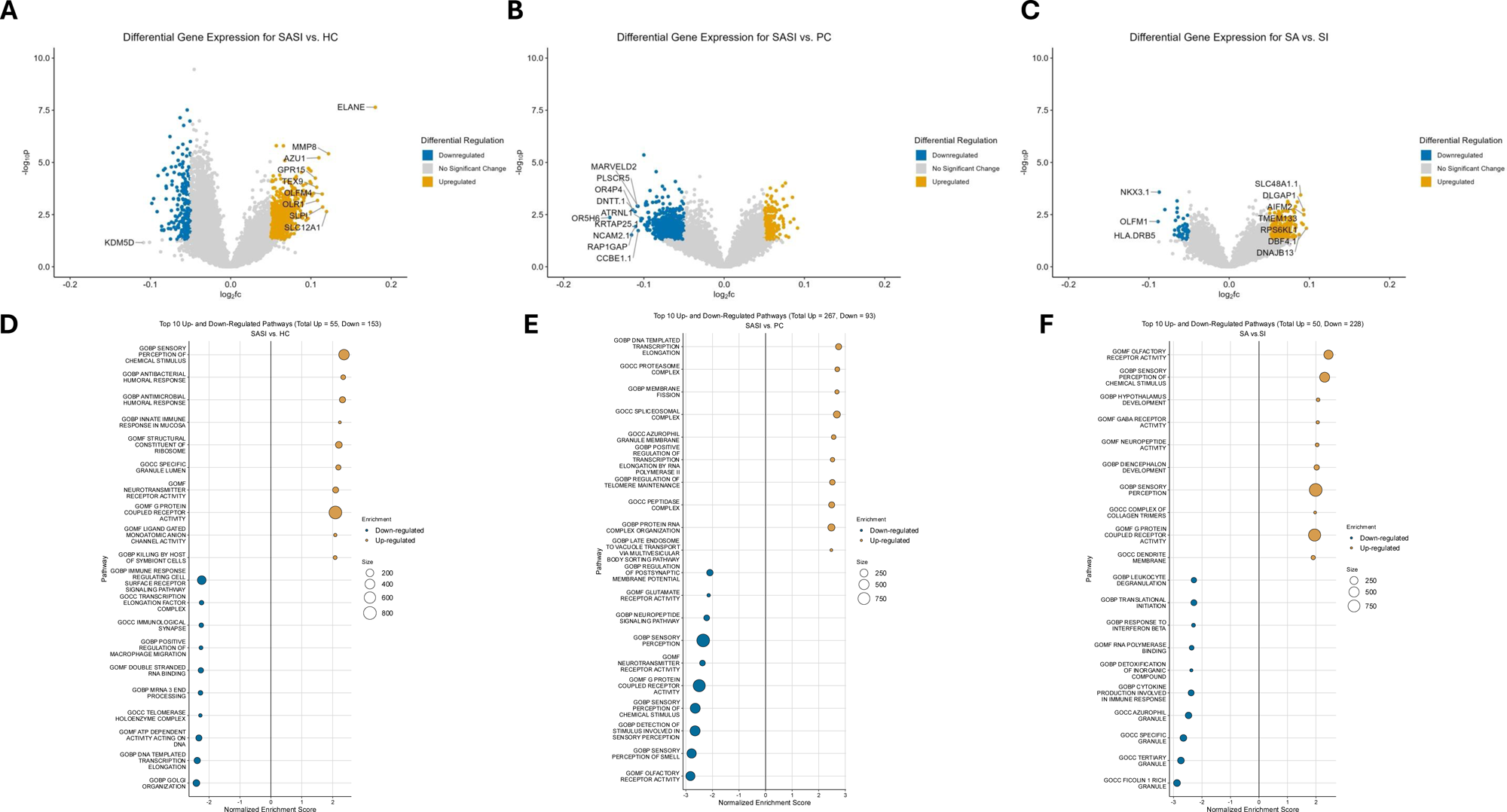
Peripheral blood transcriptomic alterations associated with suicidal thoughts and behaviors. **(A-C)** Volcano plots showing differential gene expression in blood for suicidal participants (SP) compared with healthy controls (HC) **(A)**, psychiatric comparison subjects (PC) **(B)**, and suicide attempt (SA) vs. suicidal ideation (SI) **(C)**. Genes are colored by direction of regulation (upregulated: orange: downregulated: blue, and not significant: grey). **(D-F)** Gene set enrichment analyses corresponding to each comparison panels A-C. Bubble plots display the top upregulated and downregulated pathways with normalized enrichment score on the x-axis and pathway categories on the y-axis. Bubble size reflects gene set size.

Pathway enrichment analyses revealed both distinct and convergent signatures across comparisons. In the SP vs. HC contrast (**Figure 1D**), immune-related pathways showed a bidirectional pattern of regulation. Upregulated pathways were enriched for innate and antimicrobial responses, including antibacterial defense and mucosal immunity, consistent with heightened pro-inflammatory or stress-responsive signaling. In contrast, downregulated pathways were enriched for immune-regulatory and cell-cell communication processes, such as immunological synapse organization, receptor-mediated signaling, and macrophage migration, indicating disruption of adaptive immune regulation.

In the SP versus PC comparison (**Figure 1E**), upregulated pathways were dominated by transcriptional regulation, protein synthesis, and cellular homeostasis, suggesting increased intracellular regulatory activity associated with suicidal thoughts and behaviors beyond psychiatric illness alone. Conversely, downregulated pathways were enriched for sensory perception and neuronal receptor activity, including olfactory signaling. Although detected in blood, these pathways may reflect molecular processes relevant to central sensory or perceptual systems implicated in suicide risk.

The SI vs. SA comparison revealed a distinct pattern (**Figure 1F**). Individuals with recent suicide attempt showed upregulation of pathways related to sensory processing and diencephalon development, consistent with molecular programs relevant to brain regions involved in emotional and behavioral regulation.

Downregulated pathways were again immune-related, including leukocyte degranulation, cytotoxic granule formation, and cytokine-mediated signaling, suggesting attenuated immune activity associated with suicide attempt status relative to ideation.

### Co-Expression Modules in Peripheral Blood Reveal Coordinated Immune and Metabolic Networks Associated with Suicide Risk

Because DE analyses do not capture coordinated gene activity, we applied WGCNA to identify peripheral gene co-expression modules associated with suicidal thoughts and behaviors.

In the SP vs. HC comparison, two modules were identified: Cluster 12 (8 genes; dCor=0.024; stability=0.390) and Cluster 22 (3 genes; dCor=0.025; stability=0.636; **Table 2**). Cluster 12 included inflammasome and stress-responsive genes (*CARD6, CARD17, CACNA1E, BMX, ANKRD22*), consistent with innate immune activation, whereas Cluster 22 comprised apoptosis- and immune-related genes (*BAD, ZMAT5*), suggesting altered immune homeostasis in suicidal participants relative to healthy controls.

In the SP vs. PC comparison, a single 7-gene module emerged (Cluster 8; dCor=0.027; stability=0.488). This module included immune and apoptotic regulators (*HMGN1, PSIP1, CASP16P, ZNF160*) alongside genes involved in chromatin organization and intracellular trafficking (*HIRA, EHD1*), indicating coordinated modulation of immune signaling and transcriptional regulation beyond diagnostic effects.

In the SA vs. SI comparison, a 4-gene module was identified (Cluster 10; dCor=0.017; stability=0.240), comprising *MAN2C1, MRPL51, RPL36AL,* and *RPS23*. The predominance of mitochondrial and ribosomal components implicates translational machinery and cellular energy metabolism in distinguishing suicide attempt from ideation.

To extend these findings beyond cross-sectional contrasts, we performed outcome-based WGCNA using baseline peripheral gene expression associated with subsequent suicidal behavior during follow-up. Two additional modules emerged: Cluster 41 (25 genes; dCor=0.030; stability=0.446) and Cluster 81 (7 genes; dCor=0.026; stability=0.368). Cluster 41 combined immune regulators (*IL32, IKBKB, STAT5B, HLA-DPB1*) with mitochondrial and translational genes (*NDUFB7, FLAD1, RPLP2, RPS2*), suggesting convergence of inflammatory and metabolic programs linked to future risk. Cluster 81 further implicated metabolic and ribosome-associated processes (*INSIG1, RBFA, TCP1*).

Finally, we evaluated whether genes from identified modules predicted prospective suicidal outcomes using Cox proportional hazards models over 12 months (**Supplementary Table S1**). Elevated expression of *CASP16P* (HR=1.50, 95% CI [1.19, 2.88]) and *ANKRD22* (HR=1.85, 95% CI [1.19, 2.88]) was associated with increased risk, whereas higher expression of *MRPL51*, *C20orf197*, *TBC1D30*, *ADPRH*, and *RBFA* were associated with reduced risk. Several of these transcripts also significantly predicted the broader definition of suicidal behavior, including *ANKRD22*, *CARD6*, *ZMAT5* (increased risk), and *ADPRH* (reduced risk). Together, these findings highlight coordinated peripheral gene network, particularly immune and metabolic networks, that may serve as candidate molecular markers of suicide risk.

### Attenuated Neuroimmune and Vascular Signaling in Brain Tissue from Suicide Decedents

To identify central molecular signatures associated with suicide death, we performed DE analysis in postmortem DLPFC and sgACC tissue across three contrasts: (1) suicide decedents with psychiatric illness (SP) vs. non-psychiatric comparisons (NPC), (2) SP vs. non-suicide psychiatric comparisons (NSP), and (3) NSP vs. NPC. Genes were again considered DE at a p-value < 0.05 and an absolute log_2_ fold change > 0.26.

Suicide-related comparisons yielded robust transcriptional differences in both regions. In SP vs. NPC, 773 genes were DE in the DLPFC (**Figure 2A**; 239 up, 534 down) and 1,086 in sgACC (**Figure 2B**; 455 up, 631 down). Even stronger effects were observed for SP vs. NSP, with 888 DE genes in the DLPFC (**Figure 2C**; 259 up and 629 down) and 1,304 in the sgACC (**Figure 2D**; 522 up and 782 down). Although the NSP vs. NPC contrasts also showed substantial DE signal, (DLPFC: 623 DE genes; **Figure 2E**; sgACC: 1,113 DE genes; **Figure 2F**), the larger magnitude and consistency of changes in the suicide-related contrasts suggest molecular alterations beyond psychiatric illness alone, while underscoring incomplete suicide specificity.

**Figure 2.**
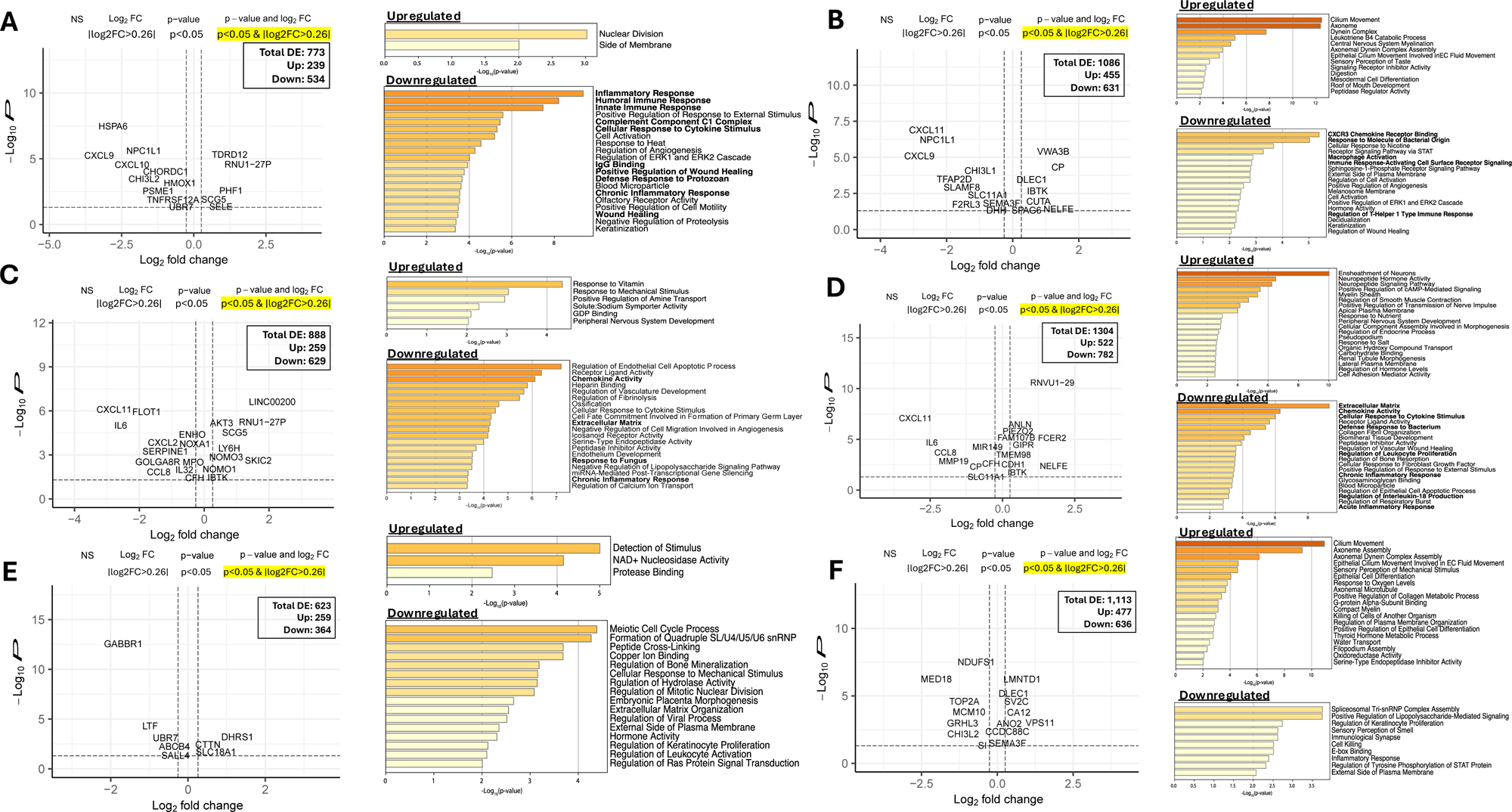
Postmortem brain transcriptomic signatures associated with suicide death. **(A-F)** Volcano plots and pathway enrichment analyses in postmortem cortical tissue from the dorsolateral prefrontal cortex (DLPFC) and subgenual anterior cingulate cortex (sgACC). Volcano plots display differential gene expression for **(A)** DLPFC, suicide vs. nonpsychiatric comparison subjects (SP vs. NPC); **(B)** sgACC, SP vs. NPC; **(C)** DLPFC SP vs. non-suicide psychiatric comparison subjects (SP vs. NSP); **(D)** sgACC, SP vs. NSP; **(E)** DLPFC NSP vs. NPC; and **(F)** sgACC, NSP vs. NPC. Points represent individual transcripts plotted by log_2_fold-change on the x-axis and -log10(p-value) on the y-axis. Genes are colored by significance thresholds with genes highlighted in red meeting our criteria of p<0.05 and |log2FC>0.26|. For each comparison, bar plots to the right show Gene Ontology pathway enrichment results for upregulated and downregulated genes, ranked by -log_10_(p-value).

Across both cortical regions, suicide decedents exhibited marked suppression of neuroimmune and vascular signaling. In the DLPFC, SP vs. NPC comparisons revealed downregulation of innate and humoral immune responses, cytokine signaling, complement activity, and wound-healing pathways (**Figure 2A**). Similar suppression was observed in SP vs. NSP, encompassing chemokine signaling, inflammatory responses, endothelial development, angiogenesis, and fibrinolysis (**Figure 2C**), indicated broad attenuation of immune and neurovascular processes. Upregulated pathways in both contrasts were comparatively sparse and reflecting intracellular and signaling processes including nuclear division and amine transport.

The sgACC showed a parallel but regionally distinct pattern. Suicide-related contrast demonstrated strong downregulation of immune signaling pathways, including macrophage activation, cytokine and STAT-mediated signaling, T-helper cell response, and vascular remodeling (**Figure 2B,D**). In contrast, upregulated pathways were enriched for ciliary function, myelination, and axonemal organization, neuropeptide signaling and neuronal ensheathment, suggesting structural and connectivity-related remodeling within this affective cortical region.

By comparison, NSP vs. NPC contrasts in both regions reflected broader psychiatric illness-associated transcriptional changes. In the DLPFC, this included modest immune suppression alongside altered sensory detection and metabolic signaling (**Figure 2E**), whereas in the sgACC, enrichment patterns included oxidative stress responses, cytoskeletal and ciliary organization, and epithelial differentiation, with concurrent downregulation of sensory signaling and RNA processing (**Figure 2F**). These findings indicate partial overlap with suicide-related changes but also highlight distinct molecular features of psychiatric illness independent of suicide.

### Brain Region–Specific Co-Expression Modules Highlight Synaptic, Immune, and Oxidative Stress Pathways in Suicide

To assess coordinated gene network disruptions associated with suicide, we applied WGCNA to postmortem DLPFC and sgACC transcriptomes. Across both regions, multiple co-expression modules were significantly associated with suicide-related contrasts (SP vs. NPC and SP vs. NSP), with additional modules emerging in psychiatric comparisons. While some biological themes were shared, several modules were region-specific, indicating anatomically distinct network-level alterations linked to suicide.

In the DLPFC, suicide-related co-expression changes were captured by multiple modules. In the SP vs. NPC comparison, Cluster 16 (**Figure 3A**; 57 genes; dCor=0.197; stability=0.172) was associated with suicide and enriched for myelination, actin cytoskeletal organization, membrane protein localization, and apoptotic signaling, suggesting disruptions in cellular structure, adhesion, and turnover. The strongest DLPFC signal emerged in the SP vs. NSP comparison where Cluster 11 (**Figure 3B**; 267 genes; dCor=0.208; stability=0.266) showed broad enrichment for gliogenesis, extracellular matrix organization, epithelial differentiation, angiogenesis, and platelet-derived growth factor signaling, consistent with altered neurovascular, developmental and wound-healing processes. In contrast, the NSP vs. NPC comparison identified a smaller module (Cluster 23; 62 genes; dCor=0.155; stability=0.134) without coherent functional enrichment, suggesting that coordinated network changes in the DLPFC are more pronounced in suicide than in psychiatric illness alone.

**Figure 3.**
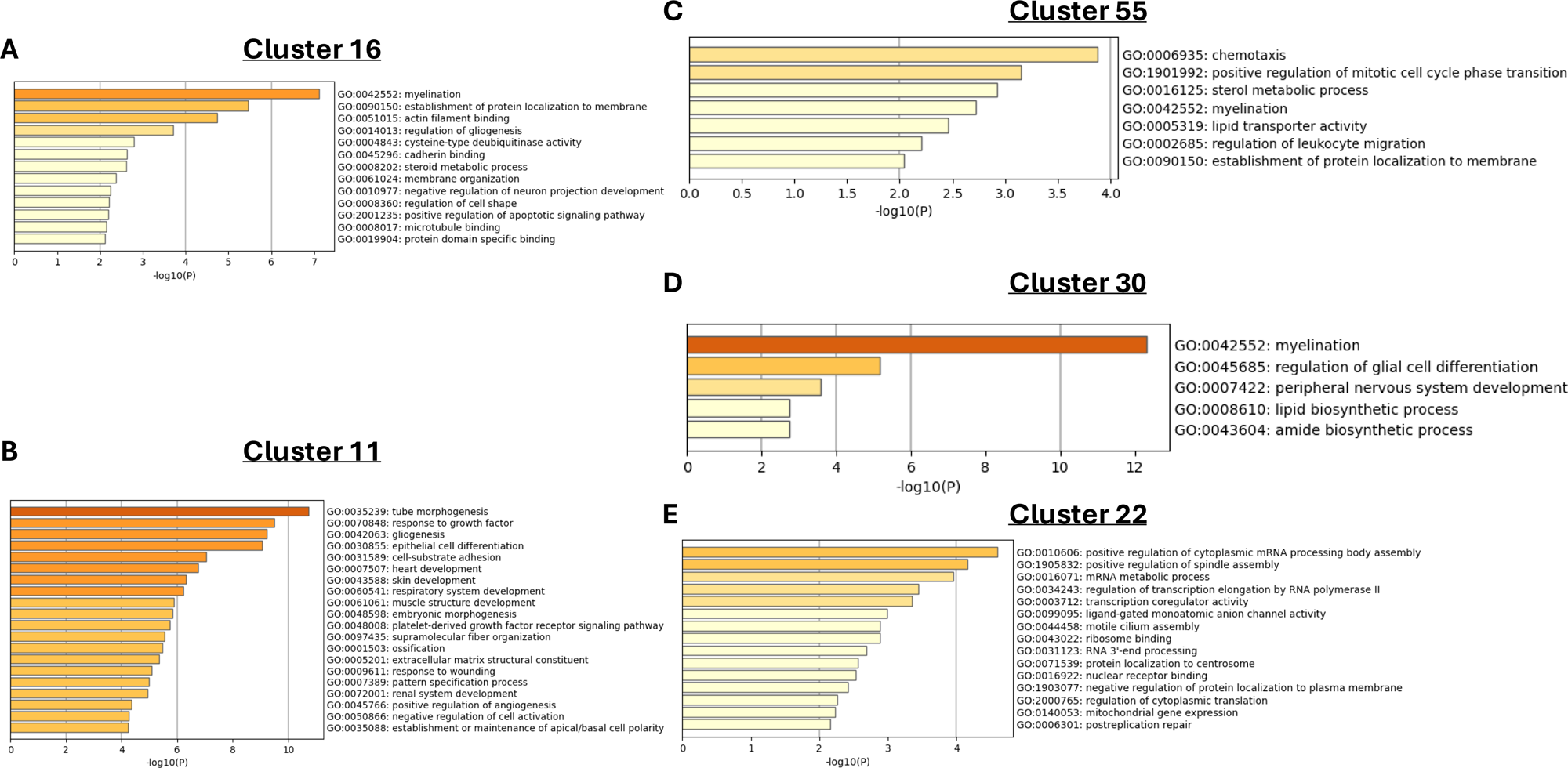
Brain region-specific co-expression modules associated with suicide. Weighted gene co-expression network analysis (WGCNA) identified suicide-associated gene modules in postmortem cortical tissue, with functional characterization of module constituents performed using pathway enrichment analyses. **(A-B)** In the DLPFC, Cluster 16 (**A**; SP vs. NPC) and Cluster 11 (**B**; SP vs. NSP) showed significant associations with suicide and were enriched for processes related to myelination, cytoskeletal organization, extracellular matrix remodeling, and developmental signaling. **(C-D)** in the sgACC, Cluster 55 (**C**; SP vs. NPC) and Cluster 30 (**D**; SP vs. NSP) were associated with suicide and showed enrichment for immune cell migration, lipid transport and metabolism, myelination and glial differentiation pathways. **(E)** Cluster 22, identified in the NSP vs. NPC comparison in sgACC as enriched for RNA processing and transcriptional regulation, reflecting molecular changes associated with psychiatric illness independent of suicide. Bars represent the top enriched Gene Ontology biological process terms ranked by -log_10_(p-value).

In the sgACC, suicide-associated modules revealed a distinct molecular profile. In SP vs. NPC, Cluster 55 (**Figure 3C**; 39 genes; dCor=0.210; stability=0.442) showed strong association with suicide and was enriched for chemotaxis, leukocyte migration, lipid transport, sterol metabolism, and myelination, indicating coordinated immune, lipid, and glial signaling alterations. In SP vs. NSP, Cluster 30 (**Figure 3D**; 34 genes; dCor=0.076; stability=0.976) was enriched for myelination, glial differentiation, and lipid biosynthesis, highlighting potential alterations in oligodendrocyte function and myelin maintenance. By contrast, the NSP vs. NPC comparison identified a large module (Cluster 22); **Figure 3E**; 298 genes; dCor=0.173; stability=0.058) enriched for RNA processing and transcriptional regulation, suggesting broader regulatory shifts associated with psychiatric illness independent of suicide.

### Cross-Region Gene Expression Overlap between Cortical Areas

RRHO analyses comparing DLPFC and sgACC transcriptomes revealed strong same-direction overlap across suicide-related and nonsuicidal psychiatric contrasts (**Supplementary Figure S1**). Concordant signal was broadly distributed across ranked gene lists and concentrated in up–up and down–down quadrants, indicating highly coordinated cortical transcriptional responses. These findings suggest that suicide-associated molecular changes are superimposed on conserved cortical regulatory programs rather than driven by large-scale inter-regional divergence.

To further characterize shared and region-specific biological processes, we performed pathway overlap analyses between DLPFC and sgACC within each contrast (**Supplementary Figure S2**). Both regions demonstrated consistent downregulation of immune, cytokine, extracellular matrix, and vascular pathways in suicide, indicating convergent suppression of neuroimmune and structural signaling. In contrast, region-specific upregulation diverged: sgACC preferentially showed enrichment for myelination, ciliary, and synaptic processes, whereas DLPFC emphasized metabolic and intracellular regulatory activity. A consolidated summary of pathway-level patterns across tissues, brain regions, and contrasts is provided in **Supplementary Figure S3**.

### Limited and Directionally Heterogenous Gene-Level Overlap between Brain and Blood

Given the strong concordance observed between cortical regions, we next examined whether suicide-associated transcriptional signals were shared between peripheral blood and brain tissue. RRHO analyses revealed contrast-dependent discordance across blood–brain comparisons, with the most prominent hotspots observed in the psychiatric diagnosis-adjusted contrasts (**Figure 4C–D**) and substantially weaker signal in comparisons involving nonpsychiatric controls (**Figure 4A–B**). Importantly, hotspots were concentrated in the top left and bottom right quadrants, indicating directionally discordant overlap: genes upregulated in brain were frequently downregulated in blood, and vice versa. These findings suggest that while overlapping gene sets are engaged across tissues, their regulation differs in direction, consistent with tissue-specific transcriptional responses rather than coordinated cross-compartment expression.

**Figure 4.**
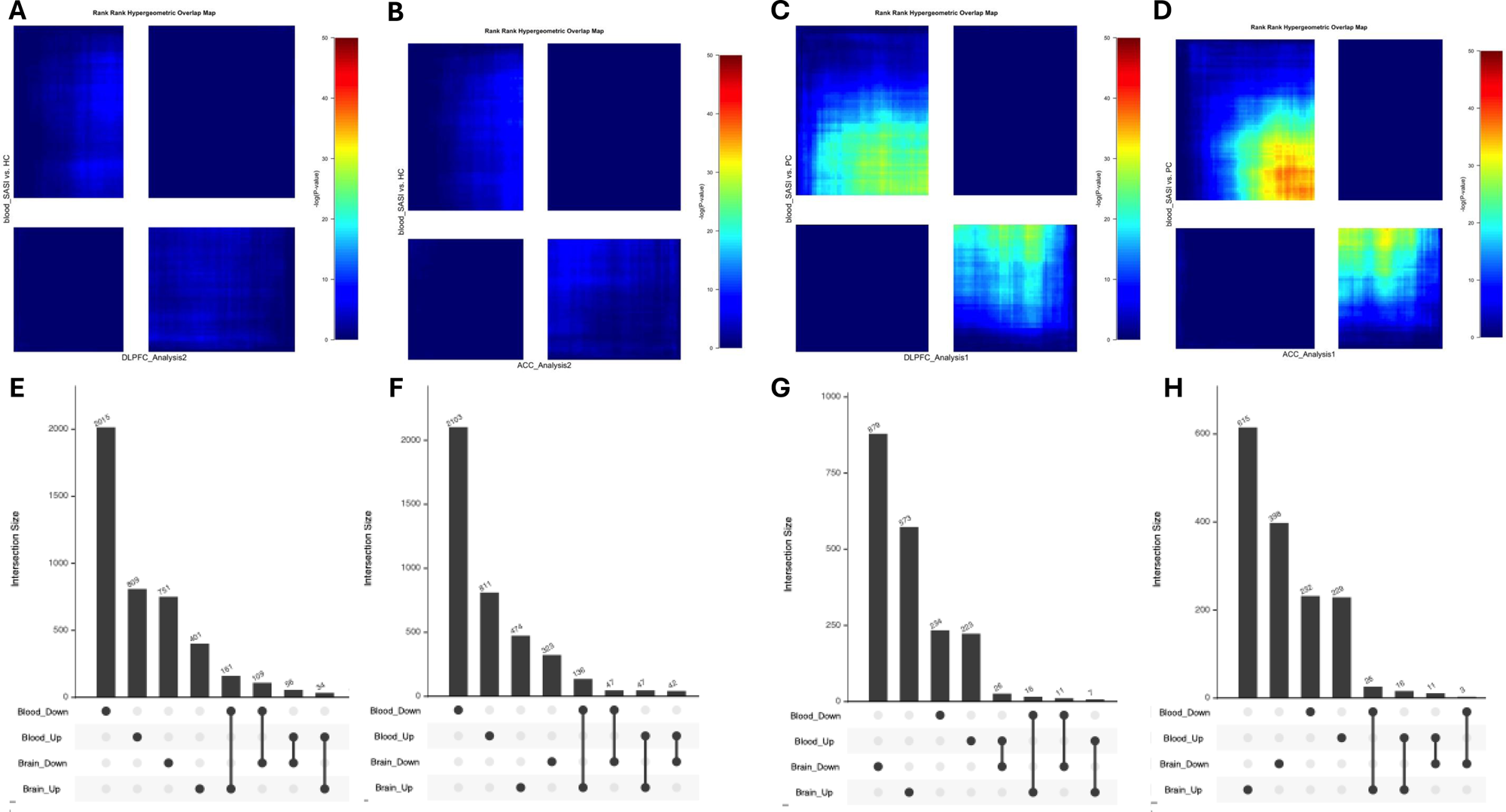
Cross-tissue gene-level overlap between peripheral and blood and cortical brain transcriptomes. **(A-D)** Rank-rank hypergeometric overlap (RRHO) heatmaps comparing ranked gene expression profiles between peripheral blood and cortical brain regions. Maps display the significance of overlap (-log_10_(p-value)) across all genes ranked by signed effect size. Concordant overlap appears in the bottom left (up-up) and top right (down-down) quadrants. Panels show comparisons between blood and DLPFC or sgACC for suicide participants vs. nonpsychiatric comparison subjects **(A-B)** and suicide participants vs. psychiatric comparison subjects **(C-D)**. **(E-H)** UpSet plots quantifying gene-level overlap between differentially expressed genes in blood and brain by direction of effect. Genes were classified as upregulated or downregulated in each tissue and intersections represent same-direction (e.g., up-up, down-down) or opposing direction (e.g. up-down, down-up) overlap). Panels correspond to blood-DLPFC **(E,G)** and blood-sgACC **(F, H)** comparisons for suicide vs. nonpsychiatric comparison subjects **(E-F)** and suicide vs. psychiatric comparison subjects **(G-H)**. Bar heights indicate the number of genes in each intersection.

To complement the rank-based RRHO analysis, we quantified overlap of threshold-defined DE genes between blood and brain using UpSet analyses stratified by direction of effect (**Figure 4E–H; Supplementary Table S2**). Across contrasts, gene-level overlap between tissues was limited relative to the total number of DE genes in each compartment and was frequently dominated by opposing-direction effects. In the SP vs. nonpsychiatric comparison, 109 genes were concordantly downregulated and 34 concordantly upregulated in both blood and DLPFC; however, substantially larger subsets exhibited discordant regulation across tissues (**Figure 4E**). A similar pattern was observed for blood–sgACC overlap in this contrast, with 47 genes concordantly upregulated and 47 concordantly downregulated, but greater discordant overlap overall (**Figure 4F**). After adjusting for psychiatric diagnosis (SP vs. psychiatric comparisons), same-direction overlap was further reduced, with only 18 concordant genes between blood and DLPFC and 19 between blood and sgACC (**Figure 4G–H**). Together, these findings indicate that cross-tissue gene-level concordance is sparse and further attenuated after accounting for psychiatric illness, with many shared genes exhibiting opposing transcriptional direction across periphery and brain.

To determine whether genes shared between blood and brain converged onto coherent biological processes, we performed pathway enrichment analyses restricted to intersecting DE genes that were regulated in the same direction across tissues (**Supplementary Figure S4**). In the SP vs. nonpsychiatric comparison, concordantly downregulated genes shared between blood and DLPFC were enriched for transcriptional and epigenetic regulation, signal transduction, immune-related pathways, developmental remodeling, and stress-response processes, whereas concordantly upregulated genes were enriched for lipid transporter activity. In contrast, same-direction overlap in SP vs. psychiatric comparisons yielded minimal enrichment, limited primarily to calcium ion binding among downregulated pathways. Similar patterns were observed for blood–sgACC overlap: in SP vs. nonpsychiatric comparisons, concordantly downregulated genes were enriched for epigenetic regulation, immune suppression, and neurodevelopmental processes, while concordantly upregulated genes were associated with antioxidant activity, translation, myeloid differentiation, and membrane-associated projections. Enrichment was again minimal in SP vs. psychiatric comparisons, with limited signal involving synapse and cell junction organization.

Taken together, RRHO and UpSet analyses indicate that gene-level overlap between peripheral blood and brain is limited, directionally discordant, and strongly context dependent. Rank-based analyses revealed structured but predominantly opposite-direction overlap across tissues, while explicit intersection of threshold-defined DE genes confirmed that shared transcriptional signals are confined to relatively small subsets and are further reduced after accounting for psychiatric diagnosis. Importantly, although cross-tissue concordance is sparse at the individual gene level, a subset of overlapping transcripts may still hold biomarker relevance. However, for mechanistic interpretation of suicide-related molecular convergence, shared biological pathways and systems-level programs provide a more coherent framework than individual genes alone.

## DISCUSSION

By integrating peripheral blood from living individuals with suicidal behavior and postmortem cortical tissue from suicide decedents, we identified convergent transcriptomic alterations involving immune, sensory, and neuronal signaling, alongside molecular signatures that were highly tissue and region-specific. In peripheral blood, suicidal thoughts and behaviors were associated with dysregulation of immune, sensory, and stress-response pathways, several of which prospectively predicted suicidal outcomes. In contrast, suicide decedents showed broad attenuation of neuroimmune and vascular signaling across both cortical regions, accompanied by region-specific upregulation: the DLPFC exhibited increased metabolic and intracellular regulatory activity, whereas the sgACC showed heightened myelination, ciliary, and synaptic-related signaling. Despite minimal gene-level overlap between tissues, shared enrichment of biological pathways suggests engagement of common systems that manifest through distinct transcriptional programs in the brain and periphery, potentially shaped by transcriptional and post-transcriptional regulatory mechanisms. Together, these findings support a coordinated yet tissue-specific molecular architecture of suicide risk.

A central contribution of this study is the explicit cross-tissue comparison of peripheral and cortical transcriptomes. Using complementary rank-based and set-based approaches, we found that gene-level overlap between blood and brain was sparse, contrast-dependent, and largely directionally discordant. Global analyses revealed modest but diffuse alignment across ranked gene lists, while set-based overlap analyses showed that only small subsets of genes exhibited concordant directionality, with most DE genes displaying tissue-specific or opposing patterns. Notably, same-direction overlap was more prominent when suicide was contrasted with nonpsychiatric comparison subjects than when psychiatric diagnoses were accounted for, suggesting that shared peripheral-central signals are attenuated after adjustment for psychiatric illness. Collectively, these findings indicate that suicide is not characterized by a conserved transcriptional signature across tissues, reinforcing the importance of pathway and systems-level approaches for interpreting cross-tissue molecular relationships.

Within this limited but consistent overlap, several genes implicated in immune regulation and transcriptional control emerged. *IFR1* and *IL6R*, key regulators of inflammatory signaling and immune homeostasis,^33,34^ were downregulated across blood and brain in suicide compared to nonpsychiatric comparison subjects, with *IFR1* showing consistent downregulation in both the DLPFC and sgACC. In parallel, shared downregulation of epigenetic regulators *SETD1A* and *CBX7*, central to chromatin-mediated transcriptional control,^35,36^ across tissues suggests disruption of transcriptional regulation that may contribute to downstream molecular vulnerability. Although neither gene has previously been associated with suicide, extensive evidence implicates *SETD1A* in schizophrenia and neurodevelopmental disorders,^37–39^ supporting the possibility that altered transcriptional regulation represents a transdiagnostic vulnerability mechanism.

In contrast, genes involved in synaptic organization and circuit-level processes showed a distinct pattern. *ROBO1* and *NPTX2*, which play established roles in axon guidance and synaptic plasticity,^40,41^ were upregulated in both blood and sgACC in suicide compared to nonpsychiatric comparison subjects. While *NPTX2* has been reported to be downregulated in cortical tissue in schizophrenia,^42,43^ its upregulation here contrasts with prior findings and aligns with evidence linking *NPTX2* to suicide attempt in genetic association studies of childhood-onset mood disorders^44^. This context-dependent pattern raises the possibility that *NPTX2* reflects suicide-related synaptic or circuit adaptations distinct from those observed in chronic psychiatric illness.

Although the genes discussed above highlight shared mechanistic themes across tissues, a subset of transcripts identified in our prospective analyses may hold greater immediate biomarker potential. Notably, *CASP16P* and *ANKRD22* were associated with increased risk of subsequent suicidal behavior, whereas *MRPL51, ADPRH*, and *RBFA* were associated with reduced risk. Several of these transcripts also participated in cross-tissue co-expression modules, underscoring their potential relevance for longitudinal risk stratification and translational validation.

Across analyses, immune-related pathways were among the most consistent signals, with peripheral blood showing mixed inflammatory and stress-related activity and both cortical regions exhibiting attenuated neuroimmune and vascular signaling. This pattern aligns with models in which chronic systemic inflammation is accompanied by insufficient or dysregulated central immune response,^45,46^ where impaired neuroimmune and glial support may compromise neuroprotection and repair. Sensory and stimulus-detection pathways also emerged across tissues, potentially reflecting altered interoceptive processing or heightened sensitivity to internal cues, mechanisms implicated in the escalation from suicidal ideation to action^47,48^.

Region-specific cortical signatures further refined these findings. In the DLPFC, upregulated metabolic and intracellular regulatory pathways may reflect compensatory demands on cognitive control systems, whereas increased myelination, ciliary, and synaptic-signaling in the sgACC suggests altered neuronal connectivity and affective integration. Given the sgACC’s role in coordinating autonomic, sensory, and emotional information, these changes may reflect disrupted processing of emotionally salient or stress-related stimuli within corticolimbic circuits.

Our results both support and clarify prior transcriptomic studies of suicide. Our results both support and refine prior transcriptomic studies of suicide. Consistent with Sun et al. (2024),^8^ we observed convergence at the level of immune-related co-expression networks across blood and brain, despite largely discordant and limited overlap at the individual gene level. Although the direction of immune involvement varies across studies, ranging from activation^49^ to attenuation,^50,51^ immune dysregulation remains one of the most recurrent themes. Phenotypic heterogeneity likely contributes to this variability; for example, Punzi et al. (2022)^10^ demonstrated that violent suicide represents a biologically distinct subtype characterized by purinergic microglial activation, highlighting the need for more granular phenotypic resolution.

Peripheral findings also converge with prior work identifying inflammatory and stress-response pathways as markers of risk^52–54^. Our longitudinal analyses extend this literature by identifying transcripts that prospectively predicted suicidal outcomes, including increased expression of *ANKRD22* and *CASP16P* and reduced expression of *ADPRH*. Although none have been previously linked to suicide, their prospective associations underscore the value of longitudinal approaches for identifying novel peripheral risk markers. Importantly, these genes map onto broader inflammatory, stress, and metabolic pathways, suggesting system-level dysregulation rather than isolated gene effects. The emergence of sensory-related pathways in blood further supports models linking altered interoception to suicide risk^55^.

Consistent with emerging models of brain-immune crosstalk, stress-induced peripheral immune activation can influence central circuits via cytokine signaling, autonomic and neuroendocrine pathways, and afferent neural inputs,^56–58^ affecting glial activity, blood–brain barrier function, and neuromodulatory tone. At the same time, correspondence between peripheral and central molecular markers is often modest at the level of individual genes, even when shared pathways are disrupted.^8,59^ Within this framework, our findings suggest coordinated engagement of shared biological systems across tissues and expressed through distinct transcriptional program.

This study has several strengths, including integration of a prospective peripheral cohort with multiregional postmortem brain data, analysis of two corticolimbic regions, and use of complementary gene-level, network-level, and pathway-level approaches.

The longitudinal design further enables assessment of future risk beyond cross-sectional associations. Limitations include phenotypic heterogeneity inherent in suicide, differences between peripheral and postmortem biologic contexts, use of bulk tissue and reliance on exploratory DE thresholds, underscoring the need for replication.

Future studies should incorporate sex- and age-stratified analyses,^60–62^ single-cell and spatial transcriptomics,^63^ and multi-omic integration^64^ to resolve cell type-specific mechanisms and regulatory hierarchies. Functional validation using iPSC-derived models^65^ and animal studies^66^ will be essential to establish causal links between transcriptomic changes, circuit dysfunction, and suicidal behavior.

In summary, this integrative analysis demonstrates that suicide is characterized not by a single, conserved molecular signature, but by coordinated disruption of shared biological systems expression through tissue-specific transcriptional program. These findings provide a foundation for developing peripheral markers that reflect central vulnerability and may ultimately inform personalized approach to suicide prevention.

## Acknowledgements

Postmortem human brain tissue was obtained from the University of Pittsburgh Brain Tissue Donation Program and the NIH NeuroBioBank at the University of Pittsburgh. We would like to thank the staff and technicians who work diligently as part of the NIH NeuroBioBank at the University of Pittsburgh. Most importantly, we thank all the family members who donated tissue of their loved ones.

---

**Supplementary Figure S1.** Cross-region rank–rank hypergeometric overlap (RRHO) between DLPFC and sgACC transcriptomes across diagnostic contrasts. Rank–rank hypergeometric overlap (RRHO) analyses were used to assess transcriptome-wide concordance between the dorsolateral prefrontal cortex (DLPFC) and subgenual anterior cingulate cortex (sgACC). (**A**) Suicide decedents with psychiatric illness (SP) vs. nonpsychiatric comparisons (NPC); (**B**) SP vs. nonsuicidal psychiatric comparisons (NSP); (**C**) NSP vs. NPC. Heatmaps display −log10 hypergeometric p-values across ranked gene lists, with warmer colors indicating stronger overlap. Enrichment in the same-direction quadrants (up–up and down–down) across contrasts indicates robust concordance of cortical transcriptional responses between regions.

**Supplementary Figure S2.** Pathway overlap analyses between DLPFC and sgACC across suicide-related contrasts. To further characterize shared and region-specific biological processes, gene ontology (GO) enrichment analyses were performed separately in the dorsolateral prefrontal cortex (DLPFC) and subgenual anterior cingulate cortex (sgACC) for each contrast. Differentially expressed genes were defined using nominal p < 0.05 and |log₂ fold change| ≥ 0.26, and genes were analyzed collectively without stratification by direction of effect. The heatmap displays −log10(P) values for enriched GO terms across DLPFC (SP vs. NPC; SP vs. NSP) and sgACC (SP vs. NPC; SP vs. NSP). Rows represent GO terms and columns represent region–contrast combinations. Hierarchical clustering was applied to both pathways and contrasts to highlight shared and region-specific enrichment patterns.

**Supplementary Figure S3.** Consolidated summary of pathway-level enrichment patterns across peripheral blood and cortical regions. Schematic overview summarizing major pathway-level enrichment patterns identified across peripheral blood, postmortem dorsolateral prefrontal cortex (DLPFC), and subgenual anterior cingulate cortex (sgACC) in suicide versus nonpsychiatric (SP vs. NPC) and suicide versus psychiatric (SP vs. NSP) comparisons. Pathways are grouped according to direction of differential expression (upregulated or downregulated) within each tissue and contrast. Categories shown reflect recurrent biological themes derived from gene ontology enrichment analyses and are intended to highlight shared and region-specific patterns across tissues.

**Supplementary Figure S4.** Pathway enrichment of same-direction intersected differentially expressed genes shared between blood and brain. To determine whether genes shared between peripheral blood and postmortem cortical tissue converged onto coherent biological processes, gene ontology (GO) enrichment analyses were performed on intersecting differentially expressed genes that were regulated in the same direction across tissues. Analyses were conducted separately for DLPFC–blood and sgACC–blood overlaps within suicide versus nonpsychiatric (SP vs. NPC) and suicide versus psychiatric (SP vs. NSP) comparisons. Upregulated and downregulated intersected gene sets were analyzed independently. Bar plots display −log10(P) values for significantly enriched GO terms. “N/A” indicates contrasts in which no significant pathway enrichment was observed.

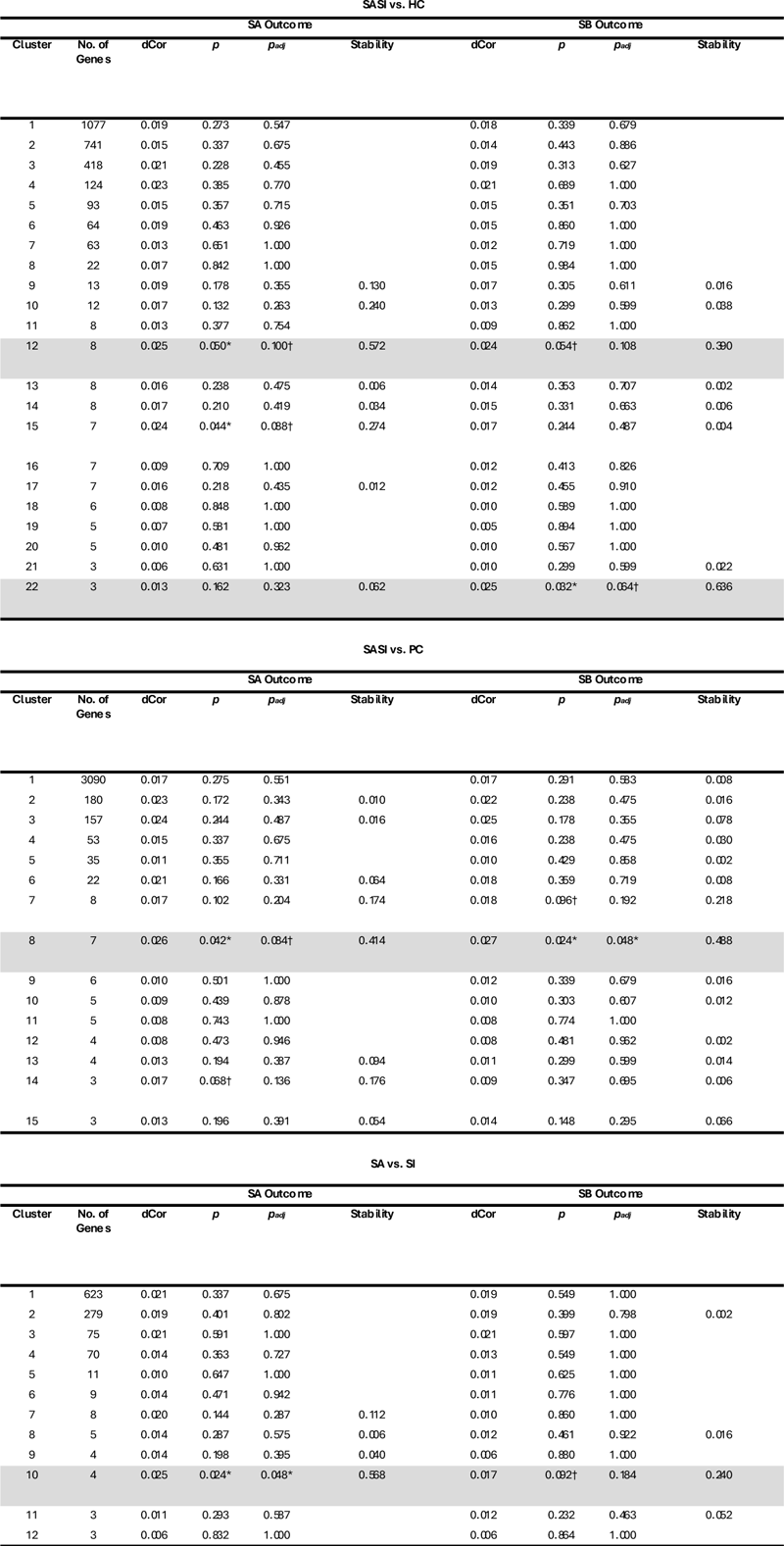

