## Supplementary Tables 1-2 for "Unraveling Tissue-Specific Molecular Signatures and Convergent Pathway Enrichments in Suicidal Behavior"

**Supplementary Table 1. Univariate Cox Models Predicting SA and SB Outcomes for Screened Clusters**

|  | **SA Outcome** | | |  | **SB Outcome** | | |
| --- | --- | --- | --- | --- | --- | --- | --- |
| **Cluster 8** | **HR (95% CI)** | ***p*** | ***p_adj_*** |  | **HR (95% CI)** | ***p*** | ***p_adj_*** |
| C22orf39; HIRA | 0.80 (0.53, 1.21) | 0.284 | 0.568 |  | 0.85 (0.63, 1.14) | 0.275 | 0.550 |
| CASP16P | 1.85 (1.19, 2.88) | 0.006** | 0.012* |  | 1.37 (1.00, 1.88) | 0.053† | 0.107 |
| EHD1 | 0.74 (0.52, 1.06) | 0.099† | 0.199 |  | 0.88 (0.66, 1.18) | 0.387 | 0.775 |
| HMGN1 | 0.72 (0.47, 1.12) | 0.142 | 0.284 |  | 0.74 (0.54, 1.00) | 0.053† | 0.106 |
| LINC00521 | 1.36 (0.96, 1.92) | 0.083† | 0.166 |  | 1.25 (0.96, 1.63) | 0.101 | 0.201 |
| PSIP1 | 0.80 (0.58, 1.10) | 0.171 | 0.341 |  | 0.82 (0.65, 1.04) | 0.095† | 0.189 |
| ZNF160-1 | 0.78 (0.53, 1.15) | 0.205 | 0.409 |  | 0.81 (0.62, 1.06) | 0.124 | 0.249 |
| **Cluster 10** | **HR (95% CI)** | ***p*** | ***p_adj_*** |  | **HR (95% CI)** | ***p*** | ***p_adj_*** |
| MAN2C1 | 1.49 (0.90, 2.45) | 0.119 | 0.238 |  | 1.25 (0.90, 1.74) | 0.186 | 0.372 |
| MRPL51 | 0.60 (0.39, 0.93) | 0.022* | 0.044* |  | 0.74 (0.55, 1.01) | 0.061† | 0.122 |
| RPL36AL | 0.78 (0.53, 1.15) | 0.213 | 0.425 |  | 0.84 (0.63, 1.11) | 0.217 | 0.434 |
| RPS23 | 0.79 (0.58, 1.08) | 0.140 | 0.279 |  | 0.86 (0.67, 1.11) | 0.245 | 0.489 |
| **Cluster 12** | **HR (95% CI)** | ***p*** | ***p_adj_*** |  | **HR (95% CI)** | ***p*** | ***p_adj_*** |
| ANKRD22 | 1.85 (1.19, 2.88) | 0.006** | 0.013* |  | 1.51 (1.11, 2.05) | 0.008** | 0.016* |
| BMX | 1.32 (0.81, 2.17) | 0.268 | 0.537 |  | 1.03 (0.73, 1.44) | 0.880 | > 0.999 |
| CACNA1E | 1.32 (0.82, 2.14) | 0.254 | 0.508 |  | 1.24 (0.89, 1.72) | 0.206 | 0.412 |
| CARD17 | 1.23 (0.78, 1.95) | 0.374 | 0.748 |  | 1.30 (0.95, 1.79) | 0.104 | 0.207 |
| CARD6 | 1.50 (0.94, 2.40) | 0.087† | 0.174 |  | 1.50 (1.08, 2.08) | 0.016* | 0.032* |
| CLEC4D | 1.43 (0.88, 2.33) | 0.150 | 0.299 |  | 1.25 (0.89, 1.74) | 0.197 | 0.393 |
| FBXL5-1 | 1.14 (0.71, 1.83) | 0.590 | > 0.999 |  | 1.05 (0.75, 1.46) | 0.784 | > 0.999 |
| KBTBD7 | 1.09 (0.67, 1.77) | 0.728 | > 0.999 |  | 1.10 (0.79, 1.54) | 0.569 | > 0.999 |
| **Cluster 22** | **HR (95% CI)** | ***p*** | ***p_adj_*** |  | **HR (95% CI)** | ***p*** | ***p_adj_*** |
| BAD | 0.90 (0.56, 1.43) | 0.646 | > 0.999 |  | 0.84 (0.61, 1.15) | 0.265 | 0.529 |
| BLOC1S1-RDH5; RP11-644F5.10 | 1.42 (0.85, 2.36) | 0.178 | 0.357 |  | 1.44 (1.02, 2.05) | 0.040* | 0.080† |
| ZMAT5 | 1.51 (0.91, 2.50) | 0.115 | 0.23 |  | 1.50 (1.06, 2.13) | 0.021* | 0.042* |
| **Cluster 41** | **HR (95% CI)** | ***p*** | ***p_adj_*** |  | **HR (95% CI)** | ***p*** | ***p_adj_*** |
| C20orf197 | 0.50 (0.31, 0.78) | 0.003** | 0.005** |  | 0.72 (0.53, 1.00) | 0.047* | 0.095† |
| FAM86C1 | 1.33 (0.86, 2.05) | 0.201 | 0.402 |  | 1.19 (0.87, 1.61) | 0.277 | 0.554 |
| FLAD1 | 1.27 (0.78, 2.07) | 0.330 | 0.66 |  | 1.07 (0.76, 1.49) | 0.700 | > 0.999 |
| FMO5 | 0.82 (0.52, 1.29) | 0.384 | 0.769 |  | 0.87 (0.63, 1.19) | 0.376 | 0.751 |
| GZMM | 1.20 (0.75, 1.90) | 0.445 | 0.889 |  | 1.11 (0.81, 1.53) | 0.505 | > 0.999 |
| HIST1H1C.1 | 1.49 (0.89, 2.49) | 0.130 | 0.26 |  | 1.34 (0.95, 1.90) | 0.093† | 0.186 |
| HLA-DPB1 | 1.30 (0.72, 2.34) | 0.383 | 0.766 |  | 1.19 (0.82, 1.72) | 0.365 | 0.73 |
| HSD17B13 | 1.23 (0.77, 1.97) | 0.394 | 0.787 |  | 1.16 (0.84, 1.60) | 0.370 | 0.741 |
| HYI | 0.83 (0.52, 1.33) | 0.449 | 0.898 |  | 0.86 (0.63, 1.19) | 0.369 | 0.738 |
| IKBKB | 0.64 (0.41, 1.00) | 0.049* | 0.098† |  | 0.73 (0.54, 1.00) | 0.050† | 0.101 |
| IL32 | 1.07 (0.65, 1.76) | 0.777 | > 0.999 |  | 0.95 (0.69, 1.32) | 0.763 | > 0.999 |
| MLLT4 | 0.85 (0.53, 1.36) | 0.499 | 0.997 |  | 0.96 (0.69, 1.33) | 0.789 | > 0.999 |
| NDUFB7 | 1.37 (0.88, 2.13) | 0.168 | 0.336 |  | 1.10 (0.80, 1.52) | 0.555 | > 0.999 |
| NKX2-5 | 0.99 (0.61, 1.61) | 0.982 | > 0.999 |  | 1.09 (0.78, 1.52) | 0.614 | > 0.999 |
| NOXRED1 | 0.97 (0.60, 1.58) | 0.905 | > 0.999 |  | 0.82 (0.58, 1.15) | 0.245 | 0.49 |
| PACS2 | 0.97 (0.61, 1.56) | 0.912 | > 0.999 |  | 0.90 (0.66, 1.23) | 0.507 | > 0.999 |
| PAFAH1B3 | 0.99 (0.61, 1.59) | 0.956 | > 0.999 |  | 0.96 (0.69, 1.33) | 0.820 | > 0.999 |
| PRM2 | 1.34 (0.87, 2.07) | 0.189 | 0.378 |  | 1.01 (0.73, 1.40) | 0.953 | > 0.999 |
| RPLP2; SNORA52 | 1.18 (0.70, 1.98) | 0.535 | > 0.999 |  | 1.13 (0.80, 1.59) | 0.503 | > 0.999 |
| RPS2; SNORA64; SNORA10 | 1.43 (0.71, 2.87) | 0.311 | 0.621 |  | 1.02 (0.73, 1.43) | 0.887 | > 0.999 |
| RRAS | 0.82 (0.51, 1.30) | 0.395 | 0.790 |  | 0.82 (0.59, 1.14) | 0.235 | 0.47 |
| STAT5B | 1.01 (0.63, 1.63) | 0.954 | > 0.999 |  | 1.11 (0.80, 1.54) | 0.521 | > 0.999 |
| TBC1D30 | 0.51 (0.30, 0.89) | 0.017* | 0.033* |  | 0.84 (0.60, 1.17) | 0.295 | 0.59 |
| ZNF511 | 0.63 (0.41, 0.97) | 0.036* | 0.072† |  | 0.72 (0.53, 0.97) | 0.032* | 0.065† |
| ZNF688 | 1.42 (0.92, 2.18) | 0.113 | 0.226 |  | 1.09 (0.80, 1.50) | 0.578 | 1 |
| **Cluster 81** | **HR (95% CI)** | ***p*** | ***p_adj_*** |  | **HR (95% CI)** | ***p*** | ***p_adj_*** |
| ADPRH | 0.56 (0.38, 0.84) | 0.005** | 0.010* |  | 0.68 (0.51, 0.91) | 0.008** | 0.017* |
| ARL15 | 1.26 (0.73, 2.18) | 0.414 | 0.828 |  | 1.12 (0.79, 1.58) | 0.527 | > 0.999 |
| FAM43A | 0.87 (0.54, 1.40) | 0.566 | > 0.999 |  | 0.83 (0.60, 1.15) | 0.256 | 0.512 |
| INSIG1 | 0.84 (0.56, 1.26) | 0.401 | 0.801 |  | 0.97 (0.71, 1.31) | 0.822 | > 0.999 |
| RBFA; RBFADN | 0.51 (0.30, 0.86) | 0.012* | 0.024* |  | 0.69 (0.49, 0.98) | 0.038* | 0.076† |
| SMIM10L1 | 0.67 (0.40, 1.11) | 0.122 | 0.244 |  | 0.71 (0.50, 1.00) | 0.052† | 0.104 |
| TCP1; SNORA20; SNORA29 | 1.00 (0.63, 1.58) | 0.989 | > 0.999 |  | 1.06 (0.76, 1.48) | 0.745 | > 0.999 |
| Note. †p < .10, *p < .05, **p < .01, ***p < .001. | | | | | | | |

**Supplementary Table 2. Same-Direction Differentially Expressed Genes Shared Between Peripheral Blood and Cortical Regions Across Suicide-Related Contrasts**

| **Gene** | **SP vs HC: Blood–DLPFC** | **SP vs HC: Blood–sgACC** | **SP vs NSP: Blood–DLPFC** | **SP vs NSP: Blood–sgACC** | **Functional class** |
| --- | --- | --- | --- | --- | --- |
| **CBX7** | ↓↓ | ↓↓ |  |  | Epigenetic |
| **CYB561A3** | ↓↓ | ↓↓ |  |  | Redox |
| **DCAF11** | ↓↓ | ↓↓ |  |  | Ubiquitin/Proteostasis |
| **DRG2** | ↓↓ | ↓↓ |  |  | Unknown/Other |
| **EPHB1** | ↓↓ | ↓↓ |  |  | Synaptic/Signaling |
| **EXD3** | ↓↓ | ↓↓ |  |  | RNA_Processing |
| **GAS8** | ↓↓ | ↓↓ |  |  | Unknown/Other |
| **GBP4** | ↓↓ | ↓↓ |  |  | Immune |
| **IFFO1** | ↓↓ | ↓↓ |  |  | Cytoskeletal |
| **IRF1** | ↓↓ | ↓↓ |  |  | Immune/Transcription |
| **PRR12** | ↓↓ | ↓↓ |  |  | Unknown/Other |
| **SETD1A** | ↓↓ | ↓↓ |  |  | Epigenetic |
| **SFI1** | ↓↓ | ↓↓ |  |  | Centrosomal |
| **SLC35C2** | ↓↓ | ↓↓ |  |  | Transporter |
| **TAF4** | ↓↓ | ↓↓ |  |  | Transcription |
| **VILL** | ↓↓ | ↓↓ |  |  | Cytoskeletal |
| **WBP1** | ↓↓ | ↓↓ |  |  | Unknown/Other |
| **ANLN** | ↑↑ | ↑↑ |  |  | Cytoskeletal |
| **GPR37** | ↑↑ | ↑↑ |  |  | GPCR/Signaling |
| **LRCH2** | ↑↑ | ↑↑ |  |  | Cytoskeletal |
| **TBC1D8B** | ↑↑ | ↑↑ |  |  | Trafficking |
| **VWA5A** | ↑↑ | ↑↑ |  |  | ECM/Adhesion |
| **ENDOD1** |  |  | ↑↑ | ↑↑ | DNA_Repair |
| **TRIP12** |  |  | ↑↑ | ↑↑ | Ubiquitin/Proteostasis |
| **CLN6** | ↓↓ |  | ↓↓ |  | Unknown/Other |
| **MTFR1L** | ↓↓ |  | ↓↓ |  | Mitochondrial |
| **SH2D3C** | ↓↓ |  | ↓↓ |  | Signaling_Adaptor |
| **SULF2** | ↓↓ |  | ↓↓ |  | ECM_Remodeling |
| **AKAP3** |  | ↑↑ |  | ↑↑ | Unknown/Other |
| **ANO7** |  | ↑↑ |  | ↑↑ | Ion_Channel |
| **HAX1** |  | ↑↑ |  | ↑↑ | Apoptosis/Stress |
| **RHBDL2** |  | ↑↑ |  | ↑↑ | Protease |
| **SLC41A2** | ↑↑ |  | ↑↑ |  | Ion_Transport |
| **TMEM125** |  | ↑↑ |  | ↑↑ | Unknown/Other |
| **ABTB1** |  | ↓↓ |  |  | Unknown/Other |
| **ACAA1** | ↓↓ |  |  |  | Metabolic |
| **ACAP3** | ↓↓ |  |  |  | Trafficking |
| **AGAP6** |  | ↓↓ |  |  | Unknown/Other |
| **AKT1** | ↓↓ |  |  |  | Kinase/Signaling |
| **APH1A** | ↓↓ |  |  |  | Unknown/Other |
| **APOBEC3C** |  | ↓↓ |  |  | Immune |
| **ARHGAP31** | ↓↓ |  |  |  | Unknown/Other |
| **ARHGAP4** | ↓↓ |  |  |  | Unknown/Other |
| **ATP2A3** | ↓↓ |  |  |  | Calcium_Transport |
| **ATXN1L** | ↓↓ |  |  |  | Unknown/Other |
| **AXIN1** | ↓↓ |  |  |  | Wnt_Signaling |
| **BANK1** |  | ↓↓ |  |  | Immune |
| **BCL9L** | ↓↓ |  |  |  | Wnt_Signaling |
| **BCR** | ↓↓ |  |  |  | Unknown/Other |
| **CASS4** |  |  | ↓↓ |  | Unknown/Other |
| **CCDC120** |  | ↓↓ |  |  | Unknown/Other |
| **CCDC61** | ↓↓ |  |  |  | Unknown/Other |
| **CD320** | ↓↓ |  |  |  | Unknown/Other |
| **CD93** | ↓↓ |  |  |  | Immune/Adhesion |
| **CDK5RAP1** |  | ↓↓ |  |  | RNA_Modification |
| **CLCN7** | ↓↓ |  |  |  | Ion_Channel |
| **CMTR1** |  | ↓↓ |  |  | RNA_Processing |
| **CNN2** | ↓↓ |  |  |  | Cytoskeletal |
| **COLQ** |  | ↓↓ |  |  | ECM/Adhesion |
| **COPS7B** | ↓↓ |  |  |  | Unknown/Other |
| **CRTC2** |  |  | ↓↓ |  | Unknown/Other |
| **CSNK1D** | ↓↓ |  |  |  | Kinase/Signaling |
| **DACH1** |  |  |  | ↓↓ | Transcription |
| **DCHS1** |  |  | ↓↓ |  | Unknown/Other |
| **DEF8** | ↓↓ |  |  |  | Trafficking |
| **DENND3** |  | ↓↓ |  |  | Trafficking |
| **DIS3L2** | ↓↓ |  |  |  | RNA_Processing |
| **DMPK** | ↓↓ |  |  |  | Kinase/Signaling |
| **E2F4** | ↓↓ |  |  |  | Transcription |
| **E4F1** | ↓↓ |  |  |  | Unknown/Other |
| **EGFL7** | ↓↓ |  |  |  | ECM/Adhesion |
| **ETS1** | ↓↓ |  |  |  | Transcription |
| **FAAP100** | ↓↓ |  |  |  | DNA_Repair |
| **FAM160B2** | ↓↓ |  |  |  | Unknown/Other |
| **FAM193B** | ↓↓ |  |  |  | Unknown/Other |
| **FBRS** | ↓↓ |  |  |  | Unknown/Other |
| **G6PD** | ↓↓ |  |  |  | Metabolic |
| **GGA1** | ↓↓ |  |  |  | Trafficking |
| **GGA2** |  | ↓↓ |  |  | Trafficking |
| **GMPR2** |  | ↓↓ |  |  | Metabolic |
| **GOLGA8B** |  | ↓↓ |  |  | Unknown/Other |
| **GUCD1** | ↓↓ |  |  |  | Unknown/Other |
| **HYAL1** | ↓↓ |  |  |  | ECM_Remodeling |
| **IL6R** | ↓↓ |  |  |  | Immune |
| **ITGB3** | ↓↓ |  |  |  | ECM/Adhesion |
| **KANK3** | ↓↓ |  |  |  | Unknown/Other |
| **KCTD15** |  |  | ↓↓ |  | Unknown/Other |
| **KDM2B** |  | ↓↓ |  |  | Epigenetic |
| **KIF21B** | ↓↓ |  |  |  | Motor/Cytoskeletal |
| **KMT2D** | ↓↓ |  |  |  | Epigenetic |
| **LEF1** | ↓↓ |  |  |  | Transcription |
| **LPAR2** |  | ↓↓ |  |  | GPCR/Signaling |
| **LRRK2** |  | ↓↓ |  |  | Kinase/Signaling |
| **MAN2B2** | ↓↓ |  |  |  | Glycosylation |
| **MAU2** | ↓↓ |  |  |  | Chromatin |
| **MEFV** | ↓↓ |  |  |  | Immune |
| **MFGE8** | ↓↓ |  |  |  | Unknown/Other |
| **MGMT** |  |  |  | ↓↓ | DNA_Repair |
| **MGRN1** | ↓↓ |  |  |  | Ubiquitin/Proteostasis |
| **MLC1** | ↓↓ |  |  |  | Unknown/Other |
| **MTHFR** | ↓↓ |  |  |  | Metabolic |
| **MTMR10** | ↓↓ |  |  |  | Unknown/Other |
| **MYLK** | ↓↓ |  |  |  | Cytoskeletal |
| **NAA60** | ↓↓ |  |  |  | Unknown/Other |
| **NADK** | ↓↓ |  |  |  | Metabolic |
| **NELFB** | ↓↓ |  |  |  | Unknown/Other |
| **NFKB2** | ↓↓ |  |  |  | Immune/Signaling |
| **NLRP1** | ↓↓ |  |  |  | Inflammasome |
| **PABPN1** | ↓↓ |  |  |  | RNA_Processing |
| **PDZD4** | ↓↓ |  |  |  | Unknown/Other |
| **PELP1** | ↓↓ |  |  |  | Unknown/Other |
| **PHC1** |  | ↓↓ |  |  | Chromatin |
| **PLAT** | ↓↓ |  |  |  | Vascular/Protease |
| **PLXDC1** |  | ↓↓ |  |  | ECM/Adhesion |
| **PPARD** | ↓↓ |  |  |  | Nuclear_Receptor |
| **PPCDC** | ↓↓ |  |  |  | Unknown/Other |
| **PPP1CA** | ↓↓ |  |  |  | Unknown/Other |
| **PRKCSH** | ↓↓ |  |  |  | Unknown/Other |
| **RAB11B** | ↓↓ |  |  |  | Trafficking |
| **RAB11FIP1** |  | ↓↓ |  |  | Trafficking |
| **RAB37** |  | ↓↓ |  |  | Trafficking |
| **RAD18** |  | ↓↓ |  |  | DNA_Repair |
| **RARA** | ↓↓ |  |  |  | Nuclear_Receptor |
| **RBM4** |  | ↓↓ |  |  | RNA_Processing |
| **RHOT2** | ↓↓ |  |  |  | Mitochondrial |
| **RNF44** | ↓↓ |  |  |  | Ubiquitin/Proteostasis |
| **SCAF1** |  |  | ↓↓ |  | RNA_Processing |
| **SETD1B** | ↓↓ |  |  |  | Epigenetic |
| **SLC12A4** | ↓↓ |  |  |  | Transporter |
| **SLC19A1** |  |  | ↓↓ |  | Transporter |
| **SLC38A10** | ↓↓ |  |  |  | Transporter |
| **SLC38A5** | ↓↓ |  |  |  | Transporter |
| **SPPL2B** | ↓↓ |  |  |  | Protease |
| **SRC** | ↓↓ |  |  |  | Kinase/Signaling |
| **STAG3L2** |  | ↓↓ |  |  | Chromatin |
| **STAT6** | ↓↓ |  |  |  | Immune/Transcription |
| **STEAP4** |  | ↓↓ |  |  | Unknown/Other |
| **SUGP1** | ↓↓ |  |  |  | RNA_Processing |
| **SYVN1** | ↓↓ |  |  |  | Ubiquitin/Proteostasis |
| **TBCD** | ↓↓ |  |  |  | Cytoskeletal |
| **TCF25** | ↓↓ |  |  |  | Unknown/Other |
| **THBD** |  |  | ↓↓ |  | Vascular/Coagulation |
| **TMEM109** | ↓↓ |  |  |  | Unknown/Other |
| **TMEM184B** | ↓↓ |  |  |  | Unknown/Other |
| **TNFAIP3** |  | ↓↓ |  |  | Immune/Regulatory |
| **TNPO2** | ↓↓ |  |  |  | Unknown/Other |
| **TNS3** | ↓↓ |  |  |  | Unknown/Other |
| **TRAF3** | ↓↓ |  |  |  | Immune/Signaling |
| **TRANK1** |  | ↓↓ |  |  | Unknown/Other |
| **TRERF1** |  | ↓↓ |  |  | Unknown/Other |
| **TRMT2A** |  | ↓↓ |  |  | RNA_Modification |
| **UBE2I** | ↓↓ |  |  |  | Ubiquitin/Proteostasis |
| **UBN1** | ↓↓ |  |  |  | Chromatin |
| **WWP2** | ↓↓ |  |  |  | Ubiquitin/Proteostasis |
| **ZNF316** | ↓↓ |  |  |  | Transcription |
| **ZNF384** |  |  |  | ↓↓ | Transcription |
| **ZNF490** |  | ↓↓ |  |  | Transcription |
| **ZNF512B** | ↓↓ |  |  |  | Transcription |
| **ZNF740** |  | ↓↓ |  |  | Transcription |
| **ZNF76** | ↓↓ |  |  |  | Transcription |
| **ZNF862** |  | ↓↓ |  |  | Transcription |
| **ATG3** |  | ↑↑ |  |  | Autophagy |
| **ATP11B** | ↑↑ |  |  |  | Transporter |
| **BCL7B** |  | ↑↑ |  |  | Chromatin |
| **C10orf88** | ↑↑ |  |  |  | Unknown/Other |
| **CABP7** |  |  |  | ↑↑ | Unknown/Other |
| **CEP57L1** | ↑↑ |  |  |  | Centrosomal |
| **CHST6** |  |  |  | ↑↑ | Glycosylation |
| **CLCA4** |  | ↑↑ |  |  | Ion_Channel |
| **CNIH1** | ↑↑ |  |  |  | Unknown/Other |
| **COMMD6** |  | ↑↑ |  |  | NFkB_Regulation |
| **CTNNA3** |  |  |  | ↑↑ | Unknown/Other |
| **DIRAS3** | ↑↑ |  |  |  | Unknown/Other |
| **DNAH17** |  | ↑↑ |  |  | Unknown/Other |
| **DNAJC21** |  |  | ↑↑ |  | Unknown/Other |
| **DNAJC22** |  | ↑↑ |  |  | Unknown/Other |
| **DPYSL3** |  | ↑↑ |  |  | Neurite/Cytoskeletal |
| **EMC1** | ↑↑ |  |  |  | Unknown/Other |
| **ESR1** |  | ↑↑ |  |  | Nuclear_Receptor |
| **FAM222A** |  |  |  | ↑↑ | Unknown/Other |
| **FANCL** | ↑↑ |  |  |  | DNA_Repair |
| **FARP1** |  |  |  | ↑↑ | Unknown/Other |
| **GAL3ST1** |  | ↑↑ |  |  | Lipid_Metabolism |
| **GDPD1** | ↑↑ |  |  |  | Unknown/Other |
| **GJB1** |  |  |  | ↑↑ | Unknown/Other |
| **GPBP1** | ↑↑ |  |  |  | Transcription/Regulation |
| **GPR3** |  | ↑↑ |  |  | GPCR/Signaling |
| **GPR68** |  | ↑↑ |  |  | GPCR/Signaling |
| **GREM2** | ↑↑ |  |  |  | Unknown/Other |
| **GSTO1** |  | ↑↑ |  |  | Detoxification |
| **KCTD3** |  | ↑↑ |  |  | Unknown/Other |
| **KLHL42** |  |  | ↑↑ |  | Ubiquitin/Proteostasis |
| **KPNA1** | ↑↑ |  |  |  | Unknown/Other |
| **LSMEM1** | ↑↑ |  |  |  | Unknown/Other |
| **MAG** |  | ↑↑ |  |  | Myelination |
| **MIER3** | ↑↑ |  |  |  | Unknown/Other |
| **MIPEP** | ↑↑ |  |  |  | Mitochondrial |
| **MRPL48** |  | ↑↑ |  |  | Mitochondrial_Ribosome |
| **MRPL57** |  | ↑↑ |  |  | Mitochondrial_Ribosome |
| **NPTX2** |  | ↑↑ |  |  | Synaptic_Plasticity |
| **NRAP** |  | ↑↑ |  |  | Cytoskeletal |
| **NUDCD1** | ↑↑ |  |  |  | Unknown/Other |
| **OSBPL6** | ↑↑ |  |  |  | Lipid_Metabolism |
| **OTUD7B** |  |  |  | ↑↑ | Ubiquitin/Proteostasis |
| **PARS2** |  | ↑↑ |  |  | Mitochondrial |
| **PCSK6** |  | ↑↑ |  |  | Protease |
| **PIRT** |  | ↑↑ |  |  | Unknown/Other |
| **PKIB** |  | ↑↑ |  |  | Signaling |
| **PLS3** |  |  |  | ↑↑ | Unknown/Other |
| **PNPLA8** |  | ↑↑ |  |  | Lipid_Metabolism |
| **PRDX4** |  | ↑↑ |  |  | Redox |
| **PRUNE2** | ↑↑ |  |  |  | Unknown/Other |
| **PSRC1** |  | ↑↑ |  |  | Unknown/Other |
| **PTCHD1** | ↑↑ |  |  |  | Unknown/Other |
| **RBBP9** | ↑↑ |  |  |  | Unknown/Other |
| **RICTOR** | ↑↑ |  |  |  | Unknown/Other |
| **ROBO1** |  | ↑↑ |  |  | Axon_Guidance |
| **RPL9** |  | ↑↑ |  |  | Unknown/Other |
| **RPS7** |  | ↑↑ |  |  | Unknown/Other |
| **RUFY3** |  |  | ↑↑ |  | Unknown/Other |
| **SEMA3E** | ↑↑ |  |  |  | Unknown/Other |
| **SFRP1** |  | ↑↑ |  |  | Wnt_Signaling |
| **SH3GL3** |  | ↑↑ |  |  | Trafficking |
| **SLC35F1** | ↑↑ |  |  |  | Unknown/Other |
| **SLC39A14** |  | ↑↑ |  |  | Transporter |
| **SMARCD3** |  | ↑↑ |  |  | Chromatin |
| **SOX10** |  | ↑↑ |  |  | Development/TF |
| **SST** |  | ↑↑ |  |  | Neuropeptide |
| **SYNPO2** | ↑↑ |  |  |  | Unknown/Other |
| **TP53TG5** |  |  |  | ↑↑ | Stress_Response |
| **TTC39A** | ↑↑ |  |  |  | Unknown/Other |
| **TXNDC17** |  | ↑↑ |  |  | Redox |
| **UBXN7** | ↑↑ |  |  |  | Unknown/Other |
| **WDR49** |  | ↑↑ |  |  | Unknown/Other |
| **XKR9** | ↑↑ |  |  |  | Unknown/Other |
| **ZBTB41** | ↑↑ |  |  |  | Unknown/Other |
| **ZDHHC9** |  | ↑↑ |  |  | Unknown/Other |
| **ZFAND1** | ↑↑ |  |  |  | Unknown/Other |
| **ZNF433** |  |  | ↑↑ |  | Transcription |
