## Supplementary Figures 1-4 for "Unraveling Tissue-Specific Molecular Signatures and Convergent Pathway Enrichments in Suicidal Behavior"

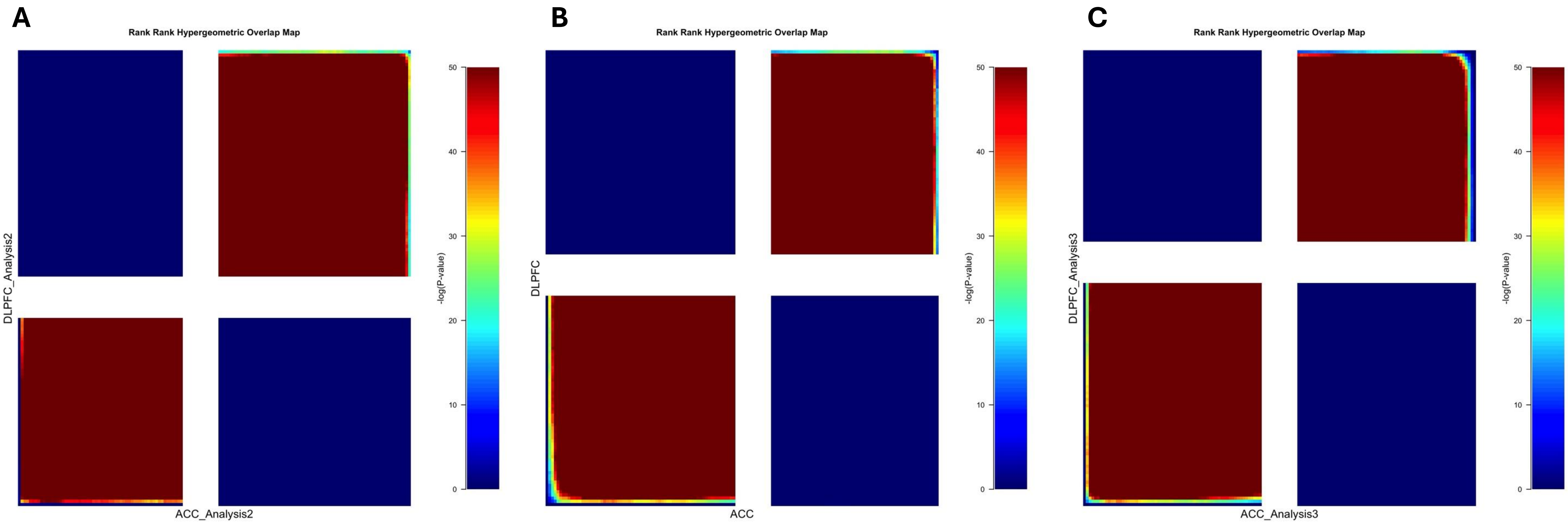

A
Rank Rank Hypergeometric Overlap Map
DLPFC_Analysis2
ACC_Analysis2
-log(P-value)
0
10
20
30
40
50
B
Rank Rank Hypergeometric Overlap Map
DLPFC
ACC
-log(P-value)
0
10
20
30
40
50
C
Rank Rank Hypergeometric Overlap Map
DLPFC_Analysis3
ACC_Analysis3
-log(P-value)
0
10
20
30
40
50

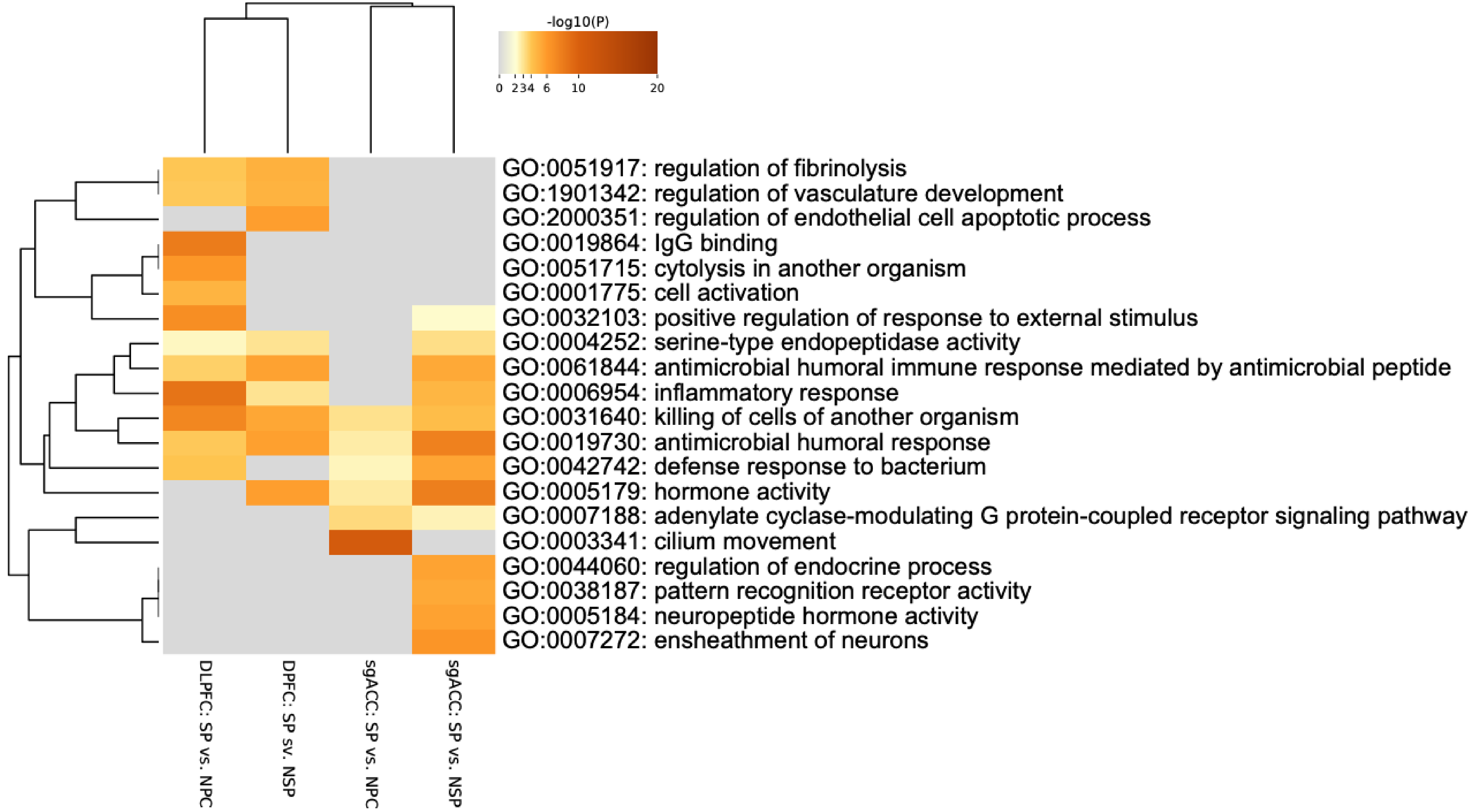
-log10(P)
0
2
3
4
6
10
20
GO:0051917: regulation of fibrinolysis
GO:1901342: regulation of vasculature development
GO:2000351: regulation of endothelial cell apoptotic process
GO:0019864: IgG binding
GO:0051715: cytolysis in another organism
GO:0001775: cell activation
GO:0032103: positive regulation of response to external stimulus
GO:0004252: serine-type endopeptidase activity
GO:0061844: antimicrobial humoral immune response mediated by antimicrobial peptide
GO:0006954: inflammatory response
GO:0031640: killing of cells of another organism
GO:0019730: antimicrobial humoral response
GO:0042742: defense response to bacterium
GO:0005179: hormone activity
GO:0007188: adenylate cyclase-modulating G protein-coupled receptor signaling pathway
GO:0003341: cilium movement
GO:0044060: regulation of endocrine process
GO:0038187: pattern recognition receptor activity
GO:0005184: neuropeptide hormone activity
GO:0007272: ensheathment of neurons
DLPFC: SP vs. NPC
DPFC: SP sv. NSP
sgACC: SP vs. NPC
sgACC: SP vs. NSP

| | Suicide vs. Non-Psychiatric Comparisons | Suicide vs. Psychiatric Comparisons |
|---|---|---|
| **Peripheral Blood** | **Upregulated**<br>• Innate immune & inflammatory signaling<br>• Antimicrobial & antibacterial responses<br>**Downregulated**<br>• Immune regulatory processes<br>• Immune-receptor mediated signaling<br>• Leukocyte migration & trafficking | **Upregulated**<br>• Transcriptional & intracellular processes<br>• Protein turnover & cellular stress responses<br>**Downregulated**<br>• Sensory perception pathways<br>• Receptor-mediated & GPCR signaling |
| **Postmortem DLPFC** | **Upregulated**<br>• Metabolic & intracellular regulation<br>**Downregulated**<br>• Immune & inflammatory signaling<br>• Cytokine & chemokine-mediated responses<br>• Vascular & extracellular matrix organization | **Upregulated**<br>• Metabolic & intracellular regulation<br>**Downregulated**<br>• Immune & inflammatory signaling<br>• Endothelial & vascular processes |
| **Postmortem sgACC** | **Upregulated**<br>• Myelination<br>• Ciliary & axonemal organization<br>• Neuronal & synaptic signaling<br>**Downregulated**<br>• Immune & inflammatory signaling<br>• Vascular & tissue repair processes | **Upregulated**<br>• Myelination<br>• Neuronal & neuromodulatory signaling<br>• MAPK-associated signaling pathways<br>**Downregulated**<br>• Chemokine & cytokine-mediated immune signaling<br>• Vascular developmental processes |

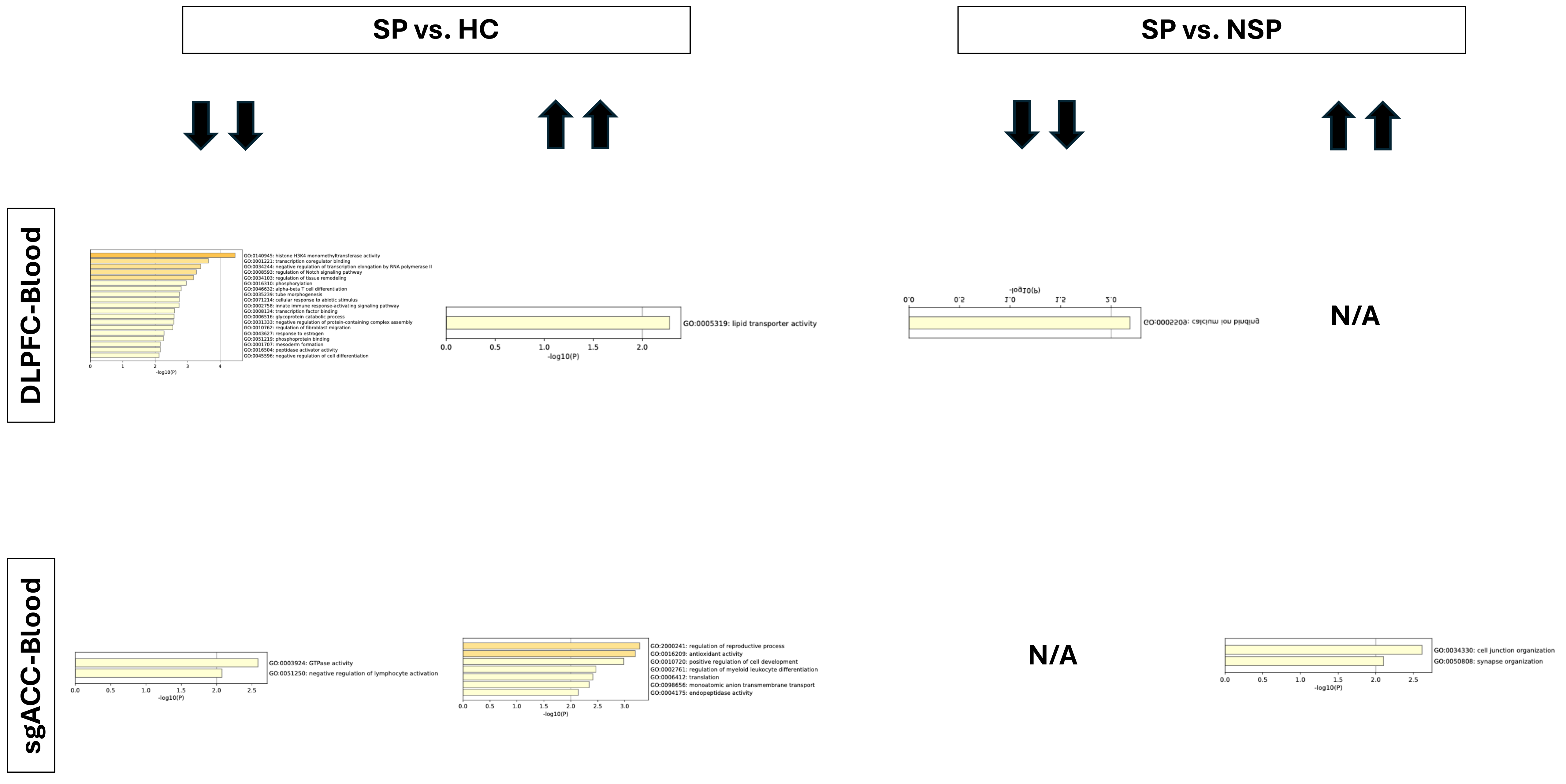
SP vs. HC
SP vs. NSP
DLPFC-Blood
sgACC-Blood
GO:0140945: histone H3K4 monomethyltransferase activity
GO:0001221: transcription coregulator binding
GO:0034244: negative regulation of transcription elongation by RNA polymerase II
GO:0008593: regulation of Notch signaling pathway
GO:0034103: regulation of tissue remodeling
GO:0016310: phosphorylation
GO:0046632: alpha-beta T cell differentiation
GO:0035239: tube morphogenesis
GO:0071214: cellular response to abiotic stimulus
GO:0002758: innate immune response-activating signaling pathway
GO:0008134: transcription factor binding
GO:0006516: glycoprotein catabolic process
GO:0031333: negative regulation of protein-containing complex assembly
GO:0010762: regulation of fibroblast migration
GO:0043627: response to estrogen
GO:0051219: phosphoprotein binding
GO:0001707: mesoderm formation
GO:0016504: peptidase activator activity
GO:0045596: negative regulation of cell differentiation
-log10(P)
GO:0005319: lipid transporter activity
-log10(P)
GO:0005509: calcium ion binding
-log10(P)
N/A
GO:0003924: GTPase activity
GO:0051250: negative regulation of lymphocyte activation
-log10(P)
GO:2000241: regulation of reproductive process
GO:0016209: antioxidant activity
GO:0010720: positive regulation of cell development
GO:0002761: regulation of myeloid leukocyte differentiation
GO:0006412: translation
GO:0098656: monoatomic anion transmembrane transport
GO:0004175: endopeptidase activity
-log10(P)
N/A
GO:0034330: cell junction organization
GO:0050808: synapse organization
-log10(P)
